# β-barrel nanopores designed for insertion into thick block copolymer membranes

**DOI:** 10.64898/2026.03.13.711555

**Authors:** Edo Vreeker, Adina Sauciuc, Fabian Grünewald, Ali Hammoudi, Giovanni Maglia

**Affiliations:** Groningen Biomolecular Sciences and Biotechnology Institute, University of Groningen, Nijenborg 7, 9747 AG, Groningen, The Netherlands; Heidelberg Institute for Theoretical Studies (HITS), Schloß-Wolfsbrunnenweg 35, 69118, Heidelberg, Germany; Interdisciplinary Center for Scientific Computing, Heidelberg University, Heidelberg, Germany; Faculty of Biosciences, Heidelberg University, Im Neuenheimer Feld 501, 69120, Heidelberg, Germany

**Keywords:** Nanopore engineering, Polymer membranes, Protein analysis, Electrophysiology, Molecular Dynamics simulations

## Abstract

Efficient integration of proteins into amphiphilic polymer membranes offers new opportunities in synthetic biology and nanotechnology. Long-term protein reconstitution into artificial membranes remains challenging due to a lack of stabilising protein-membrane interactions found in native lipid bilayers. Here, we redesigned the transmembrane region of a CytK-4D β-barrel nanopore for stable insertion into 3.5–6.6 nm thick PBD-PEO (poly(1,2-butadiene)-b-poly(ethylene oxide)) bilayers. PBD-PEO membranes offer high mechanical and chemical stability and low electrical noise, but the thick membrane hinders anchoring of biological nanopores. By systematically investigating the elongation of the β-barrel, we engineered nanopore constructs suitable for PBD_11_PEO_8_ and PBD_22_PEO_14_ membranes. Efficient insertions were observed by adding amino acids that stabilised the transmembrane β-barrel structure and enhanced anchoring of the nanopore into the membrane. Molecular dynamics simulations and single-molecule assays revealed that nanopores folded naturally into PBD-PEO bilayers, enabling successful detection of cyclodextrins and translocation of polypeptides and full-length proteins. Our study offers important lessons for the reconstitution of membrane proteins into artificial membranes. Moreover, these highly robust nanopore-membrane interfaces can be readily integrated into biosensing devices, enabling peptide and protein analysis directly from complex solutions.

## Introduction

Nanopores enable identification of biomolecules at the single-molecule level.^[1–3]^ The incorporation of nanopores into microelectromechanical systems may lead to new types of biosensors capable of analysing biological samples in real-time. A substantial bottleneck in the development of these devices is the fragility of the lipid membranes in which nanopores are conventionally reconstituted.^[4,5]^ Lipid bilayers are prone to rupture when exposed to external perturbations or challenging experimental conditions. Alternatively, amphiphilic polymers may be used as membranes, which can demonstrate higher mechanical, chemical and electrical stability.^[6–9]^ The applicability of robust polymer membranes for portable nanopore devices is exemplified by the successful commercialisation of portable flow cells developed by Oxford Nanopore Technologies for the sequencing of nucleotides.^[10,11]^

Several amphiphilic polymers have been investigated for membrane protein reconstitution. Two commercially available amphiphilic polymers include the triblock copolymer PMOXA-PDMS-PMOXA (poly(2-methyloxazoline-b-dimethylsiloxane-b-2-methyloxazoline)) and diblock copolymer PBD-PEO (poly(1,2-butadiene)-b-poly(ethylene oxide)).^[12–14]^ The electrical properties of several nanopores inserted in these polymer membranes have been reported.^[15–19]^ PBD-PEO show great promise for nanopore sensing, being resistant to high electrical potentials (∼420 mV for PBD_11_PEO_8_, and ∼540 mV PBD_22_PEO_14_)^[19]^, high concentrations of human serum (∼25% serum)^[19]^, and exposure to high concentrations of denaturing salts (guanidinium chloride, GuHCl).^[18]^

Five β-barrel nanopores have been successfully inserted into PBD_11_PEO_8_ bilayers (hydrophobic thickness *d* ≈ 3.5 nm^[20]^): MspA, ɑ-hemolysin, aerolysin, lysenin and CytK.^[18,19]^ However, these nanopores poorly anchor into PBD_11_PEO_8_ membranes, resulting in short-term membrane exits.^[19]^ Furthermore, while β-barrel nanopores ɑ-hemolysin and CytK inserted into PBD_22_PEO_14_ (*d* = 6.6 nm^[21]^) membranes, 10^3^ higher nanopore concentrations were required and, crucially, unstable current baselines prevented nanopore measurements. Other tested β-barrel nanopores (MspA, aerolysin and lysenin) could not be characterised in PBD_22_PEO_14_, due to short-term membrane exits (MspA and aerolysin) or no observed membrane insertion (lysenin).

Here we have engineered 13 new nanopore constructs containing 0.7–3.5 nm transmembrane domain (TMD) and hydrophilic extensions. Long-term reconstitution into PBD-PEO membranes was observed with nanopores containing up to 2.8 nm extensions. Extensions of both the hydrophilic and hydrophobic region of the β-barrel domain were required. Constructs yielding favourable electrical properties were identified and allowed the detection of γ-cyclodextrins and translocation of both disordered model polypeptides and full-length proteins. Molecular dynamics simulations revealed that the PBD-PEO membrane thins around the β-barrel region of the nanopore. Furthermore, the hydrophilic PEO region of the membrane most likely influences the conductivity of the nanopore by partially obstructing the nanopore entry and by interacting with potassium ions. These nanopore constructs embedded in robust polymer membranes are highly promising for applications in biosensing, for example to enable protein fingerprinting directly from biological samples.

## Results and Discussion

### CytK 4D in PDB_11_PEO_8_ and PBD_22_PEO_14_ membranes

PDB_11_PEO_8_ and PBD_22_PEO_14_ membranes have a hydrophobic core (*d)* of 3.5 and 6.6 nm, respectively, and provide enhanced chemical, mechanical and electrical stability compared to phospholipid membranes, such as DPhPC (*d* = 2.8 nm).^[22]^ Despite their favourable properties, the incorporation of nanopores in such membranes is challenging and not always possible. For example, we presented the CytK-4D nanopore, an engineered CytK nanopore with four aspartate residues equally spaced in the β-barrel region for enhanced electroosmotic flow, enabling translocation of unravelled proteins.^[23]^ Insertion of CytK-4D into robust PBD-PEO bilayers instead of fragile lipid bilayers would greatly boost biosensing and sequencing applications. However, residence times of CytK-4D into PBD_11_PEO_8_ membranes are extremely short, with nanopores frequently observed to exit shortly after membrane insertion (Figure S1). This is in contrast to wild-type CytK^[19]^, suggesting an enhanced destabilising effect by the introduced residues K128D, Q145D, S151D and K155D. When CytK-4D was tested for reconstitution into PBD_22_PEO_14_, no insertion was observed, despite using 10^5^⨉ higher concentrations than required for DPhPC, and testing inserting potentials up to +400 mV. In short, CytK-4D nanopores are incompatible with PBD-PEO bilayers.

We hypothesised that the instability of CytK-4D in PBD-PEO bilayers may be largely driven by the mismatch between the hydrophobic thickness of PBD-PEO membranes and the transmembrane domain of the nanopore, which evolved to match the hydrophobic thickness of natural lipid membranes. The TM region of CytK includes 12 amino acids between V119 and V131 and between P136 and W148 on each strand, spanning ∼4.2 nm in length, considering typical extensions of 0.35 nm per amino acid^[24–26]^. Since the hydrophobic core (*d)* of DPhPC is 2.8 nm^[22]^, the lipid bilayer and the amino acid spacing likely remodel to align their respective hydrophobic domains (**Figure 1A**).

**Figure 1.**
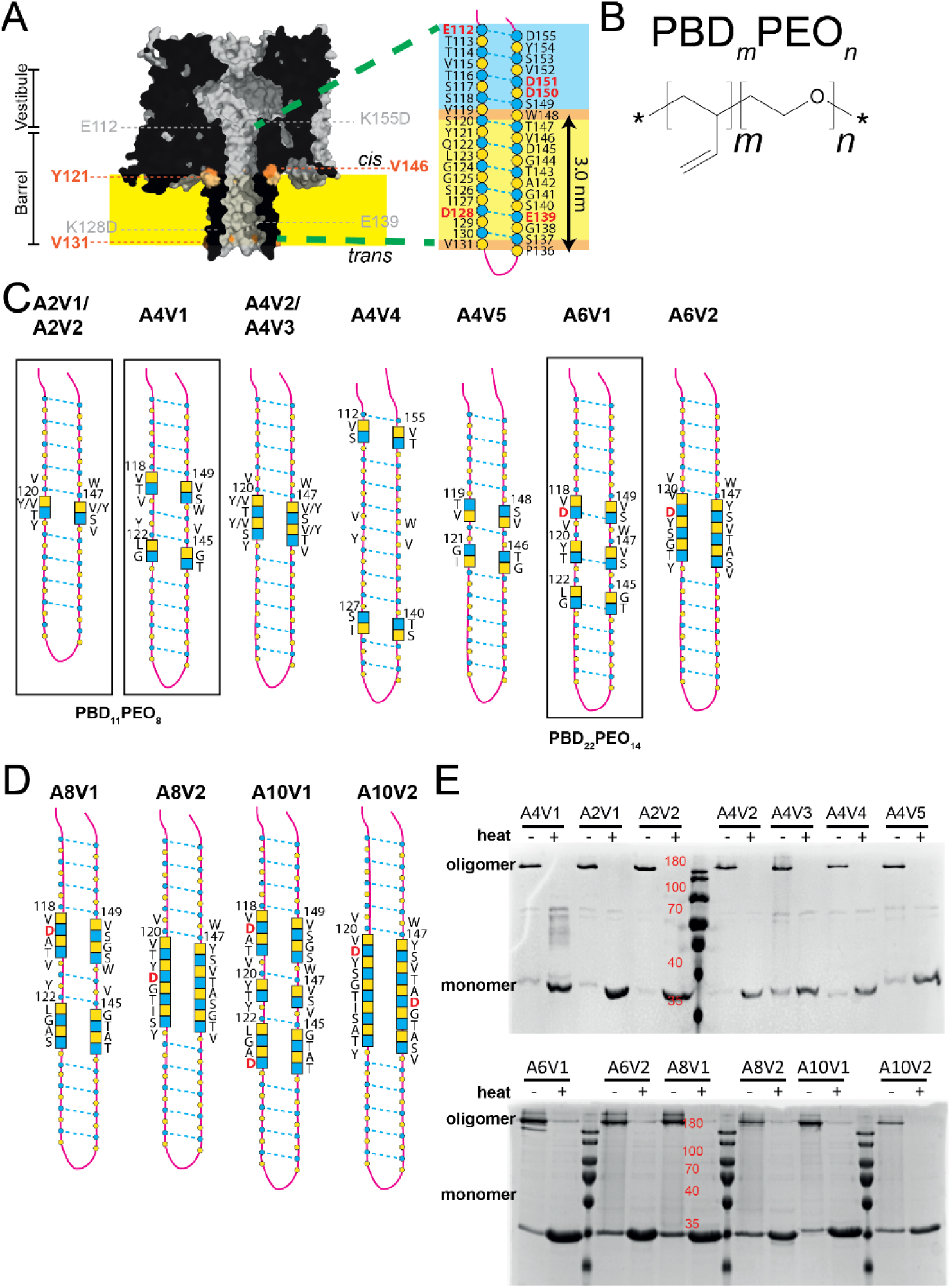
Barrel extension of the CytK-4D nanopore. A) Surface representation of a cut-through CytK-4D nanopore, where in orange the positions of residues located at the lipid bilayer-water interface (yellow, DPhPC) are indicated. Zoom-in: description of the β-barrel of the CytK-4D nanopore showing the hydrophobic region of the lipid bilayer in yellow, the region of interaction of the lipid headgroups (orange) and the soluble region (blue). B) Chemical structure of the amphiphilic polymer PBD-PEO (poly(1,2-butadiene)-b-poly(ethylene oxide)). C) Nanopore designs where the barrel is extended by two, four or six amino acids. D) Depiction of the extension of eight and ten amino acids. The amino acids in the original 4D pore are depicted as circles, and the added amino acids as squares; the residues facing the lumen are coloured in blue, while the ones facing the membrane are coloured in yellow. E) SDS-PAGE of the extended constructs. In the absence of heat, the oligomer band above 180 kDa (expected Mw of ∼248-259 kDa) is present. Upon heating the sample, this high Mw band disappears, while the monomer band of 35.5-37 kDa becomes more prominent, demonstrating the dissociation of the SDS-stable, heat-labile oligomers.

To approach the hydrophobic thickness of PBD_11_PEO_8_ and PBD_22_PEO_14_ membranes (chemical structure in **Figure 1B**), we constructed several versions of CytK-4D nanopores, where each β-sheet was elongated with two, four, six, eight and ten amino acids. The nanopore TMD was thus elongated by 0.7 – 3.5 nm, not considering the possible changes in the tilt in the β-barrel within the membrane. Several designs were tested to optimise the nanopore-membrane interaction, keeping in mind the crucial role of alternating polar and non-polar residues in beta-strands.^[27]^

In ꞵ-barrel nanopores certain amino acids (such as threonine and serine) are commonly favoured as building blocks (Table S1). Tyr is another commonly favoured residue within membrane proteins, whose position in the TMD is often found at the membrane-water interface to enhance the anchoring of membrane proteins.^[28]^ For this reason, Tyr residues have also been included in the *de novo* design of nanopores.^[29]^ Gly residues are known to stabilise ꞵ-barrel configurations in both biological and *de novo* nanopores^[29–31]^ by increasing the ꞵ-sheet structure^[29]^ or reducing the strain that could arise from steric clashes along the side chain^[31]^.

New constructs were designed inspired by the amino acid composition within the transmembrane barrel of CytK (**Figure 1 C,D**). This barrel is rich in Thr and Ser residues, but also contains multiple valine (Val) residues, making these amino acids appealing candidates to add to the TMD. Furthermore, we also considered tyrosine (Tyr) for some designs (**Figure 1 C, D**), which is naturally present in the hydrophobic belt within the barrel. Multiple insertion sites were chosen, leading to a variety of new nanopore constructs. All insertions (∇) were on both strands of the barrel to ensure symmetric extensions. To specify individual constructs, it was chosen to use the following nomenclature: AxVy, where ‘x’ specifies the number of extra amino acid (A) pairs per strand added to the CytK nanopore and ‘y’ indicates the y^th^ version (V).

To maintain the symmetry around the middle residues, Ser120-Tyr121 and Val146-Thr147 on opposite sides of the β-strand, for the first two extended constructs two amino acids (extension of ∼0.7 nm) were inserted between Ser120 and Tyr121 and between Thr147 and Val146, yielding the A2V1 and A2V2 constructs (∇2 pores, **Figure 1C, D**). For the addition of four amino acids (∇4 pores, extension of ∼1.4 nm) 5 constructs were designed following different strategies to maximise our chances of forming stable extended nanopores. In two ∇4 constructs the four amino acids were placed in the middle of the barrel (A4V2 and A4V3). One construct was prepared by inserting two residues at each extremity (E112-D155 and I127-S140 pair, A4V4), extending both the transmembrane domain and hydrophilic domain of the barrel. In two constructs, insertions were made above and below the central residues of the aromatic region [between the S118-S149 and Q122-D145 pairs (A4V1); and the V119-W148 and Y121-V146 pairs, (A4V5). All these constructs were highly expressed in *E. coli* SoluBL21(DE3). Although most of the protein remained in the insoluble fraction (Figure S2), purification with 0.02 % DDM resulted in the formation of SDS-stable, heat-labile oligomers for all two and four AA extensions (**Figure 1E**).

To elongate the hydrophobic region even further, we then tested constructs with β-sheets extended by six (∇6 pores,∼2.1 nm), eight (∇8, ∼2.8 nm) and ten (∇10, ∼3.5 nm) amino acids at both sides. A6V2, A8V2, A10V2 constructs were made by inserting the amino acids between the Ser120-Tyr121 and Thr147-Val146 pairs in the middle of the β-barrel aromatic region. To minimise the risk that the nanopore would not fold, we made additional constructs based on combinations of the A2V1 and A4V1, in which the β-barrel is elongated at different positions with respect to the *cis* and *trans* entries (A6V1, A8V1 and A10V1). These longer constructs also contained added Gly residues, to stabilise ꞵ-barrel configurations ^[29–31]^. Moreover, one or two Asp residues were included to improve the solubility of these longer constructs, provided that the distance between Asp residues could be maintained above 1 nm, in order to minimise negative impacts on EOF.^[23]^ Similarly to their shorter counterparts, all these constructs were well expressed, although mostly in the insoluble fraction (Figure S2). However, as observed with the previous constructs, they formed the characteristic CytK oligomers, which dissociated in the presence of heat (**Figure 1E**). An overview of the amino acid sequences for the transmembrane domains of all constructs is also provided in Table S2.

### Electrical characterisation of extended barrels in PBD-PEO membranes

Nanopore constructs were tested for their functionality by adding pre-formed oligomers to the buffer in the experimental chamber. To achieve consistent membrane insertion within 10 minutes, 10⨉ dilutions up to <1 µL of undiluted sample were added to the chamber, depending on the insertion activity of a particular construct into its respective membrane (PBD_11_PEO_8_ or PBD_22_PEO_14_).

For each reconstituted nanopore we measured (1 M KCl, 15 mM TRIS, pH 7.5, N>3 experiments): the open pore current (*I*_0_, ±75 mV); the root mean square (RMS) *I*_0_ fluctuation over time [(*I*_RMS_^[32]^, where larger values of *I*_RMS_ are indicative of higher electrical noise or a larger degree of nanopore gating (transient closure of the nanopore)] and *R*_sym_ (the ratio of I_0_ at +75 mV and −75 mV, which indicates the symmetry of the current-voltage relationship – the latter parameter typically deviates from unity for highly selective pores).

Nanopore constructs with up to 8 amino acids added per strand consistently inserted into both DPhPC and PBD-PEO bilayers (PBD_11_PEO_8_ for constructs with 2-4 extra amino acids per strand and PBD_22_PEO_14_ for constructs with 6-10 extra amino acids per strand). Importantly, and in contrast to the original CytK-4D nanopore, all the new constructs presented long-term stability in PBD-PEO membranes. Each nanopore construct produced unique current signals both in DPhPC and their corresponding PBD-PEO membrane (**Table 1**, Figure S3, S4). A2V1 (**Figure 2A**), A2V2 and A4V1 reconstituted in PBD_11_PEO_8_ produced favourable current properties under negative voltage, the typical working potential with CytK-4D in DPhPC. These nanopores all produced open pore current signals compatible with unobstructed nanopores, i.e. stable current baselines, low-noise electrical signals and limited nanopore gating. Other nanopore constructs reconstituted in PBD_11_PEO_8_ were found to lack at least one of these essential properties. A6V1 reconstituted in PBD_22_PEO_14_ was the only nanopore construct yielding the abovementioned favourable current properties in the thickest polymer membrane (**Figure 2A**, S3, S4, **Table 1**). The current voltage-dependencies of CytK-4D and all extended nanopores became strongly asymmetric when reconstituted in PBD-PEO bilayers, whereas insertion into DPhPC resulted in more linear current-voltage relationships (**Table 1**, Figure S5). Several extended nanopores showed inverted current-voltage dependencies in PBD-PEO relative to CytK-4D (i.e., the ionic currents at positive applied potential are higher than at negative applied potential). Interestingly, whereas A4V1, A4V2 and A4V4 followed a similar trend of CytK-4D, A4V3 and A4V5 yielded inverted current-voltage relationships, while all constructs contained the same number of additional amino acids. The difference is particularly remarkable between A4V2 and A4V3, since both nanopore constructs use the same insertion sites, and only differ by the locations of inserted Val and Tyr residues being reversed. Extended nanopores containing 6 or 8 added residues per strand (characterised in PBD_22_PEO_14_) all showed an inverted current-voltage relationship relative to CytK-4D. For the A6V1 construct it was confirmed that this behaviour was also observed in the thinner PBD_11_PEO_8_ bilayers (data not included). These observations suggest that the conductance behaviour of these nanopores is a complex phenomenon, being highly sensitive to the location and character of residues, the nature of the surrounding membrane, and likely dependent on local conformations of the nanopore-membrane complex.

**Table 1.** Open pore current values for 11 nanopore constructs in PBD-PEO membranes.

| Construct | Membrane | $\langle I_0 \rangle$<br>(-75 mV), pA | $\langle I_0 \rangle$<br>(+75 mV), pA | $\langle I_{RMS} \rangle$<br>(-75 mV), pA | $\langle I_{RMS} \rangle$<br>(+75mV), pA | $R_{sym}$ |
| --- | --- | --- | --- | --- | --- | --- |
| CytK-4D | DPhPC | $-114.8 \pm 7.5$ | $106.0 \pm 2.4$ | $1.4 \pm 0.3$ | $6.6 \pm 6.6$ | 0.92 |
| A2V1 | PBD<sub>11</sub>PEO<sub>8</sub> | $-76.1 \pm 2.5$ | $33.1 \pm 2.8$ | $1.2 \pm 0.1$ | $1.1 \pm 0.1$ | 0.43 |
| A2V2 | PBD<sub>11</sub>PEO<sub>8</sub> | $-79.9 \pm 0.1$ | $47.3 \pm 1.4$ | $1.3 \pm 0.1$ | $1.2 \pm 0.1$ | 0.59 |
| A4V1 | PBD<sub>11</sub>PEO<sub>8</sub> | $-81.2 \pm 3.0$ | $54.9 \pm 3.8$ | $1.2 \pm 0.1$ | $1.4 \pm 0.1$ | 0.68 |
| A4V2 | PBD<sub>11</sub>PEO<sub>8</sub> | $-77.7 \pm 0.9$ | $56.8 \pm 2.6$ | $2.1 \pm 1.0$ | $2.7 \pm 1.1$ | 0.73 |
| A4V3 | PBD<sub>11</sub>PEO<sub>8</sub> | $-51.2 \pm 1.9$ | $70.1 \pm 5.7$ | $1.5 \pm 0.1$ | $9.6 \pm 11.5$ | 1.37 |
| A4V4 | PBD<sub>11</sub>PEO<sub>8</sub> | $-81.4 \pm 2.5$ | $54.0 \pm 2.2$ | $1.2 \pm 0.1$ | $1.4 \pm 0.1$ | 0.65 |
| A4V5 | PBD<sub>11</sub>PEO<sub>8</sub> | $-65.9 \pm 9.7$ | $87.3 \pm 4.9$ | $1.6 \pm 0.1$ | $3.6 \pm 1.9$ | 1.32 |
| A6V1 | PBD<sub>22</sub>PEO<sub>14</sub> | $-28.7 \pm 3.8$ | $66.6 \pm 4.5$ | $1.0 \pm 0.1$ | $1.0 \pm 0.1$ | 2.32 |
| A6V2 | PBD<sub>22</sub>PEO<sub>14</sub> | $-34.3 \pm 6.6$ | $69.9 \pm 4.6$ | $2.1 \pm 0.8$ | $3.5 \pm 2.6$ | 2.04 |
| A8V1 | PBD<sub>22</sub>PEO<sub>14</sub> | $-28.9 \pm 10.1$ | $64.8 \pm 2.2$ | $1.2 \pm 0.2$ | $1.2 \pm 0.2$ | 2.24 |
| A8V2 | PBD<sub>22</sub>PEO<sub>14</sub> | $-27.2 \pm 7.5$ | $60.8 \pm 9.0$ | $1.1 \pm 0.1$ | $1.1 \pm 0.1$ | 2.24 |

$\langle I_0 \rangle$ : average open pore current values of nanopore constructs in a corresponding membrane. $\langle I_{RMS} \rangle$ : average root mean square current values, reflecting the noise of a nanopore construct in a particular membrane. $R_{sym}$ : ratio of $\langle I_0 \rangle$ at +75 mV over $|\langle I_0 \rangle|$ at -75 mV. $I_{RMS}$ and $I_0$ are averaged values obtained from 3 individual measurements (N = 3). Buffer conditions: 1 M KCl, 15 mM TRIS, pH 7.5. 10 kHz sample frequency and 2 kHz low-pass Bessel filter was used for measuring.

**Figure 2:**
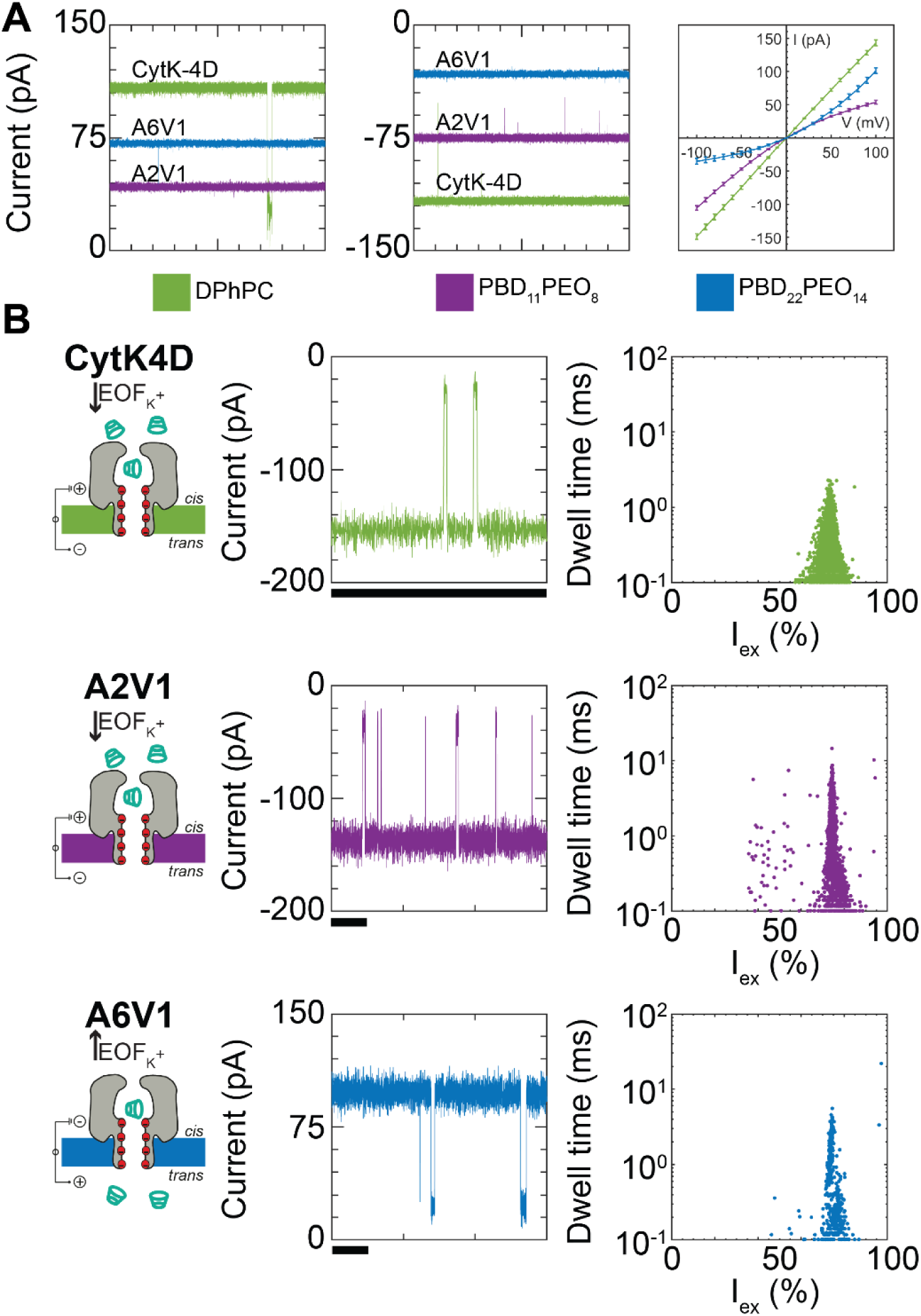
Current properties of CytK-4D, A2V1 and A6V1 and subsequent γ-cyclodextrin detection. A) Overview current properties CytK-4D (in DPhPC, green) A2V1 (in PBD_11_PEO_8_, in purple) and A6V1 (in PBD_22_PEO_14_, in blue) at +75 mV and -75 mV (left and middle panel). Buffer: 1 M KCl, 15 mM TRIS, pH 7.5. In the right panel corresponding representative IV-curves are shown. B) Single-molecule γ-cyclodextrin (γ-CD, 5 µM) detection with CytK-4D (in DPhPC, green), A2V1 (in PBD_11_PEO_8_, purple) and A6V1 (in PBD_22_PEO_14_, blue) under -100 mV (CytK-4D and A2V1) or +100 mV (A6V1). Left panels illustrate direction of the dominant electro-osmotic forces and location of γ-CD molecules (*cis* for CytK-4D and A2V1, *trans* for A6V1). Middle panels show representative blockade levels of γ-CD interacting with the nanopore. Scale bars represent 25 ms. Right panels: representative scatter plots of dwell time versus excluded current (*I*_ex_). A distinctive clustering of data points is observed for each nanopore construct, revealing that cyclodextrin molecules undergo specific interactions with each construct, thus suggesting stable barrel structures. Events were recorded using 50 kHz sampling frequency and 10 kHz low-pass Bessel filter.

### Nanopore barrel assessment with cyclodextrins

To verify that the elongated nanopores assembled as β-barrel structures in PBD-PEO membranes, similarly as CytK in DPhPC bilayers, we tested the sensing of γ-cyclodextrins (γ-CD) molecules with elongated nanopores yielding favourable current properties. Cyclodextrins interact with specific sites in the beta-barrel domain of the nanopore, and are known to yield characteristic current blockades in ɑ-hemolysin, a homolog of CytK.^[33,34]^ For A2V1, A2V2 and A4V1 in PBD_11_PEO_8_, analytes were added to the *cis*-chamber while applying a negative voltage to work with the direction of the electroosmotic flow. Since A6V1 showed an inverse current-voltage relationship compared to CytK-4D (both in PBD_22_PEO_14_ and PBD_11_PEO_8_), a positive voltage was used for single-molecule measurements, requiring analytes to be added to the *trans*-chamber to work with the direction of the EOF.

The addition of 5 µM γ-CD (*cis*) to CytK-4D in DPhPC (−100 mV, *cis*) yielded characteristic blockade levels (**Figure 2B**, S6). Events taken from three independent measurements (recording time 60 s) were used to obtain the average dwell-time and excluded current, *I*_ex_ (**Table 2**, *I*_ex_ is defined as: 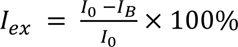, where *I*_B_ is the blocked current value during an event). The resulting scatter plot of dwell time *vs*. *I*_ex_ yielded an observable cluster (**Figure 2B**, S7). The same measurements with A2V1, A2V2 and A4V1 in PBD_11_PEO_8_, and A6V1 in PBD_22_PEO_14_ (−100 mV, γ-CD in *trans*) also yielded characteristic blockade levels and scatter plots (**Figure 2B**, S6, S7). The dwell times and *I*_ex_ increased as the β-barrel domain increased (**Table 2**). Although each construct might have slightly different tilt or folds most likely, the dwell times reflect the increased resistance of the γ-CD in longer barrels. Of note, dwell times events measured with A6V1 in PBD_22_PEO_14_ were shorter compared to the events measured with nanopore constructs in PBD_11_PEO_8_ despite the longer β-barrel domain. This effect is likely due to the *trans* to *cis* direction of the γ-CD transport, which might lead to a different interaction with the membrane or the nanopore.

**Table 2.** γ-cyclodextrin detection by CytK-4D (DPhPC, *cis*), A2V1, A2V2 and A4V1 in PBD_11_PEO_8_, (*cis*) and A6V1 in PBD_22_PEO_14_ (*trans*). Data taken from 60 s measurements (*N* = 3).

| Construct | Membrane | Voltage | Chamber | n events | Dwell time (ms) | $I_{ex}$ (%) |
| --- | --- | --- | --- | --- | --- | --- |
| CytK-4D | DPhPC | −100 mV | *cis* | >2694 | $0.23 \pm 0.06$ | $75.7 \pm 3.2$ |
| A2V1 | PBD<sub>11</sub>PEO<sub>8</sub> | −100 mV | *cis* | >1678 | $1.25 \pm 0.09$ | $74.8 \pm 0.5$ |
| A2V2 | PBD<sub>11</sub>PEO<sub>8</sub> | −100 mV | *cis* | >1649 | $2.21 \pm 0.11$ | $78.9 \pm 0.9$ |
| A4V1 | PBD<sub>11</sub>PEO<sub>8</sub> | −100 mV | *cis* | >957 | $4.00 \pm 0.15$ | $81.2 \pm 0.2$ |
| A6V1 | PBD<sub>22</sub>PEO<sub>14</sub> | +100 mV | *trans* | >329 | $0.89 \pm 0.03$ | $77.0 \pm 1.8$ |

### Molecular dynamics simulations in PBD-PEO membranes

To elucidate the possible origins of the altered ionic currents observed by the different nanopore constructs and provide a molecular interpretation of the system, we performed all-atom molecular dynamics simulations. We focused on two constructs that yielded favourable current properties in their respective polymer membranes: A2V1 in PBD_11_PEO_8_ and A6V1 in PBD_22_PEO_14_. The MD simulations showed that for both nanopores, the engineered aromatic region was shorter than the membrane’s thickness, with the pore’s *trans* exit piercing only the polymer’s hydrophobic core (**Figure 3 A, B**). In both cases, and especially for A6V1 in PBD_22_PEO_14_, the membrane thins and acquires a substantial curvature to accommodate the protein **(Figure 3 A, B**). For both nanopores, at the *trans* side the hydrophilic PEO part is located near and above the nanopore entry, interacting with the ions passing across the nanopore. The longer PEO moiety in PBD_22_PEO_14_ covered the *trans* side of the nanopore to a larger extent than A2V1 in PBD_11_PEO_8_ membranes (**Figure 3A, B**). The presence of the PEO polymers near the *trans* entry of the nanopore might help explaining the altered conductivity of the nanopores in PBD-PEO compared to lipidic membranes.

**Figure 3:**
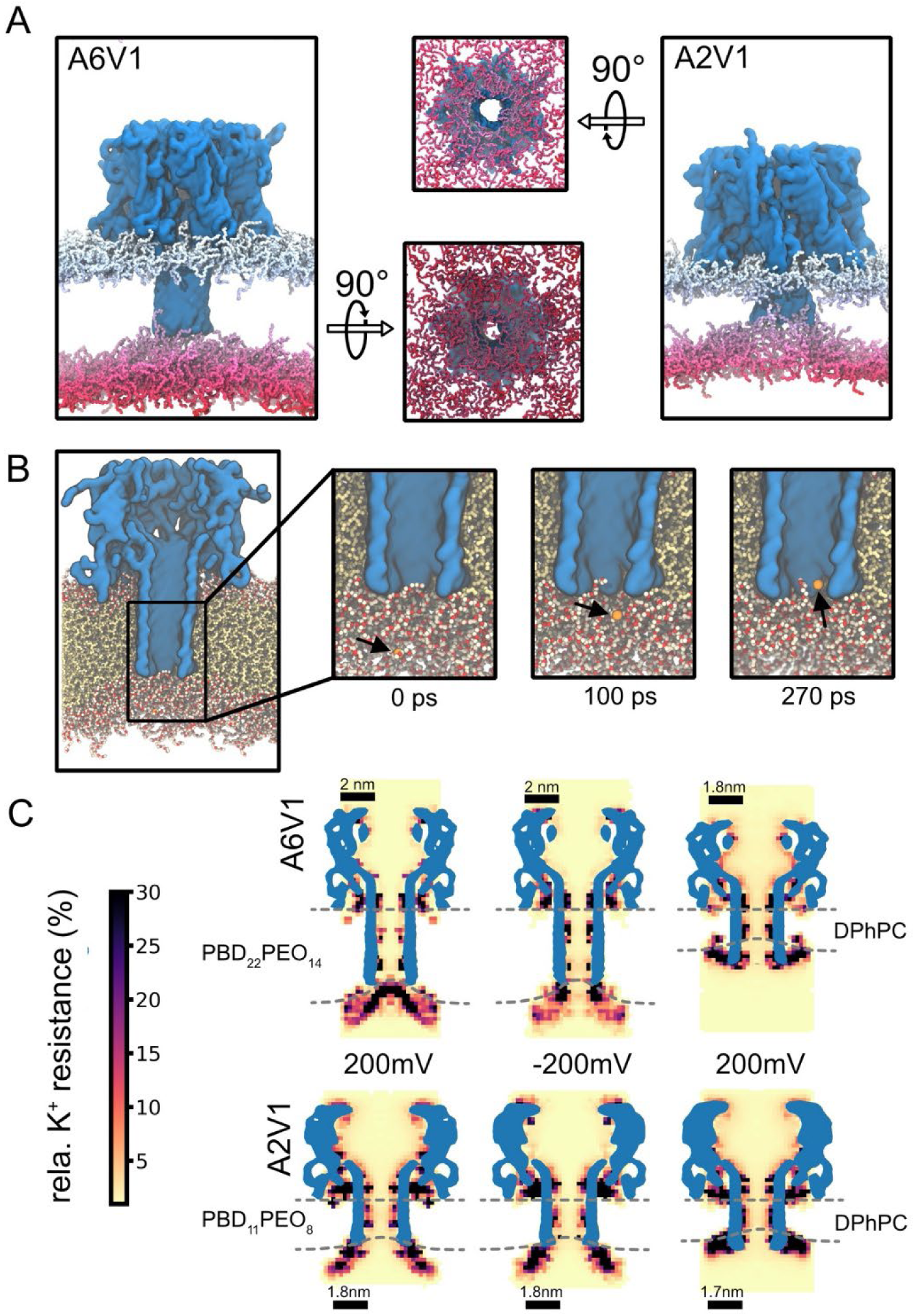
All-atom molecular dynamics simulation of pore constructs in polymer membranes. A) Snapshot of the A6V1 pore in PBD_22_PEO_14_ membranes (left) and A2V1 pore in PBD_11_PEO_8_ membranes (right, upper leaflet blue; lower leaflet red) under +200mV applied field at the end of the simulation. Water, ions, and the hydrophobic core of the membrane are omitted. B) Snapshot of A6V1 in PBD_22_PEO_14_ membranes at the end of a 250ns simulation. The nanopore surface is shown in blue, the hydrophobic polymer in yellow, and PEG in red/white. Water and ions are omitted for clarity. The zoom-in shows a potassium ion (orange) moving into the pore obstructed by the hydrophilic PEO layer at the bottom of the pore. C) Relative potassium resistance for the A6V1 and A2V1 constructs obtained from MD simulations in polymer and lipid-based membranes as indicated. The hydrophobic polymer membrane core (PBD) is indicated by the dashed lines as well as the hydrophobic core of the DPhPC membrane. Note that the A6V1 pore appears shorter in the DPhPC membrane (top row; third column) due to the *trans* end being less ordered and thus not within the density cutoff (Figure S10, S11).

To better understand the membrane’s influence on potassium-ion motion, we computed the relative potassium resistance on a grid (**Figure 3C**), which measures how much slower potassium ions move relative to the bulk potassium ion movement. For both nanopores in their respective membranes, potassium movement was slowed down at the trans entry, independently of the field direction (Figure S8, S9). According with the experimental data, the magnitude of this effect is greater for the A6V1 in PBD_22_PEO_14_ compared to the A2V1 in PBD_11_PEO_8_. In comparison, in the reference systems using a DPhPC lipid bilayer membrane, potassium motion was significantly less impaired (**Figure 3C**). Therefore, MD simulations showed that the reduced current through the nanopore embedded into PBDPEO membranes is likely caused by the occupancy of PEO moieties inside the nanopore.

Of note, when A6V1 was introduced into DPhPC membranes, the length of the nanopore was significantly shorter than in PBD_22_PEO_14_ membranes (**Figure 3C**). This is due to the *trans* end of the pore becoming more disordered to fit within the shorter DPhPC membranes (Figure S10). This structural difference might contribute to the observed reduction in conductivity when A6V1 is embedded in the polymer membrane compared to DPhPC membranes, as longer pores show a higher resistance. By contrast, the A2V1 variant did not show a significant structural difference when embedded in a short polymer membrane compared to DPhPC.

It is also worth noticing that two observations in MD simulation disagreed with experimental data. We found that for the A6V1 pore in PBD_22_PEO_14_ membranes, potassium motion was affected by the field direction (**Figure 3C**), however, in the opposite direction as observed in experiments. Further, A2V1 conductivity remained unchanged in PBD_11_PEO_8_ and DPhPC membranes, despite a change in ionic current that was observed experimentally. These observed discrepancies may arise from limitations in the applied force field or from structural differences (e.g., locally different protein folding) that are not accounted for in the protocol used to generate the initial structure.

In general, the effect of the polymer membrane on the nanopore conductance is likely to be affected by both the length of the PEO moiety, which is considerably shorter in PBD_11_PEO_8_ compared to PBD_22_PEO_14_, and by the length of the β-barrel region. The long A6V1 nanopore in PBD_22_PEO_14_ showed the most compromised conductance, compared to shorter nanopores in PBD_11_PEO_8_ or DPhPC membranes. All-atom MD simulations suggest that the PEO of the polymer membrane affects conduction at the exit of the nanopores by interacting with potassium ions. This effect adds to the increased resistivity induced by longer β-barrel domains, and might alter the interaction with analytes and the nanopore. Interestingly, despite longer designed β-barrel domains, in the DPhPC bilayers the elongated pores become more disordered and effectively shorter due to the large hydrophobic mismatch. Both effects provide a mechanistic molecular explanation for the reduction in conductivity observed for the longer nanopore constructs in artificial membranes.

### Peptide and protein detection with A2V1, A2V2, A4V1 in PBD_11_PEO_8_ and A6V1 in PBD_22_PEO_14_

An important application of nanopores is the unidirectional transport of unstructured proteins across nanopores^[23,35]^, which is only observed with nanopores with a strong EOF. Therefore, it is important that the EOF in the new nanopore constructs is retained after barrel elongation. Note that as for the experiments with γ-CD, analytes were added to *cis* for A2V1, A2V2 and A4V1 (under negative voltage, in PBD_11_PEO_8_) and to *trans* for A6V1 (under positive voltage, in PBD_22_PEO_14_).

We used the 140 AA-long model polypeptide tzatziki, with an unfolded structure and relatively high negative charge density (-7e every 100 amino acids).^[23]^ Events were characterised by short dwell times, high excluded currents and a flattened electrical signature (see **Table 3** for dwell time and *I*_ex_ values at −160 mV, N = 3) above a threshold potential of −100 mV.

**Table 3.** Tzatziki detection by CytK-4D (DPhPC, *cis*), A2V1, A2V2 and A4V1 in PBD_11_PEO_8_, (*cis*) and A6V1 in PBD_22_PEO_14_ (*trans*). Data taken from 60 s measurements (*N* = 3).

| Construct | Membrane | Threshold potential | Detection potential | Side | #events | Dwell time (ms) | $I_{ex}$ (%) |
| --- | --- | --- | --- | --- | --- | --- | --- |
| CytK-4D<sup>[23]</sup> | DPhPC | $-100$ mV | $-160$ mV | *cis* | N.A. | $0.66 \pm 0.06$ | $88.0 \pm 2.0$ |
| A2V1 | PBD<sub>11</sub>PEO<sub>8</sub> | $-120$ mV | $-160$ mV | *cis* | $>707$ | $3.91 \pm 1.01$ | $95.0 \pm 0.2$ |
| A2V2 | PBD<sub>11</sub>PEO<sub>8</sub> | $-140$ mV | $-160$ mV | *cis* | $>982$ | $2.74 \pm 0.33$ | $91.2 \pm 0.2$ |
| A4V1 | PBD<sub>11</sub>PEO<sub>8</sub> | $-140$ mV | $-160$ mV | *cis* | $>856$ | $4.80 \pm 2.00$ | $91.9 \pm 1.8$ |
| A6V1 | PBD<sub>22</sub>PEO<sub>14</sub> | $+160$ mV | $+160$ mV | *trans* | $>246$ | $5.65 \pm 1.69$ | $93.2 \pm 0.9$ |

Characteristic blockade signals were observed for all extended nanopores with the dwell time of events decreasing upon increasing voltage, which is a hallmark for translocation (**Figure 4A**, S12‒S15). The resulting dwell time values were consistently longer for all extended nanopores relative to CytK-4D in DPhPC under the same applied voltage (**Table 3**). Threshold potentials and *I*_ex_ were higher relative to CytK-4D in DPhPC (∼20-40 mV and 3-5%pt, respectively, N> 3, **Table 3**). These outcomes most likely reflect the reduced EOF from the lower ionic currents produced by the elongated nanopores.

**Figure 4:**
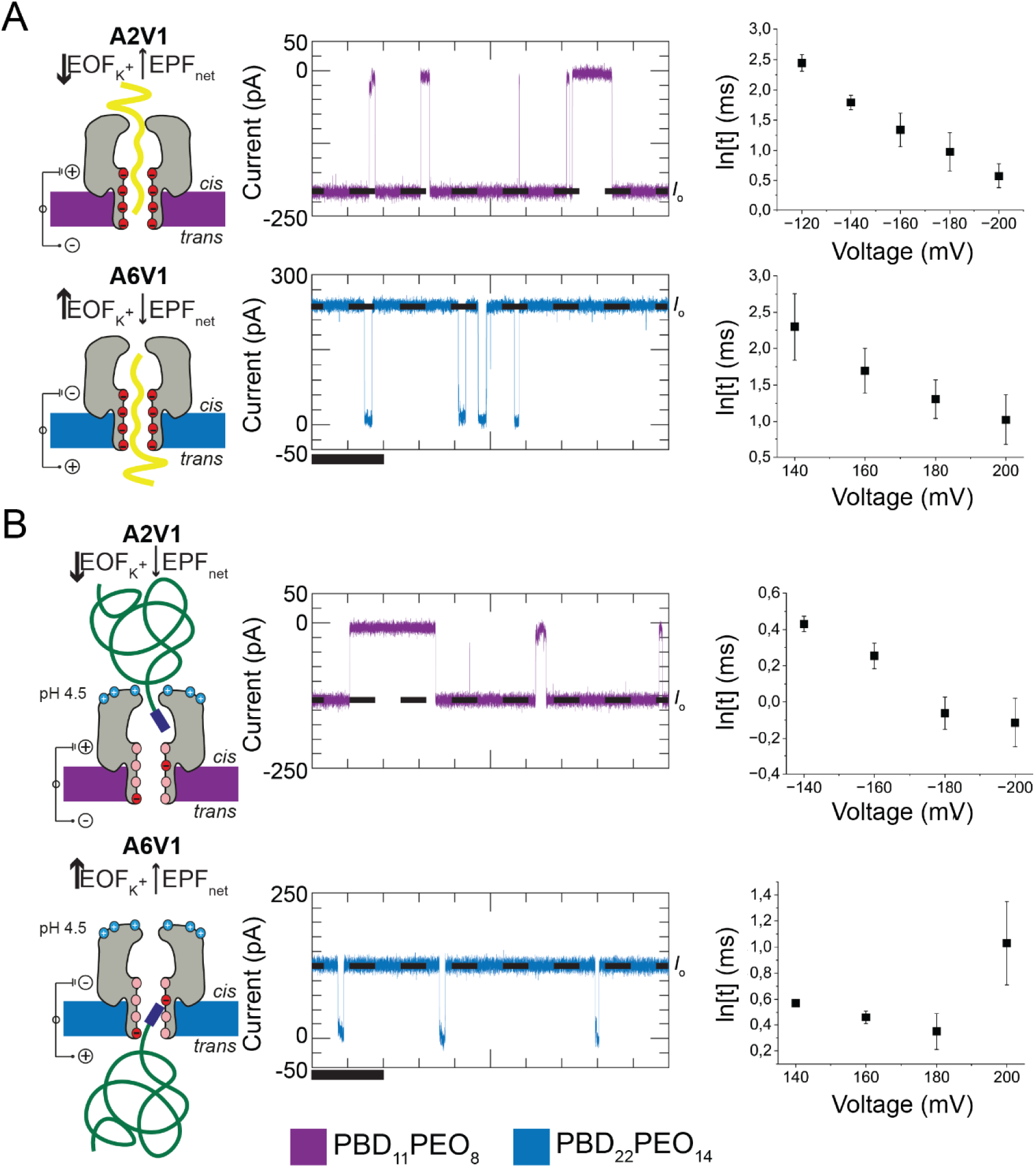
Polypeptide and full-length protein translocation with A2V1 (PBD_11_PEO_8_) and A6V1 (PBD_22_PEO_14_). On the left the direction of electroosmotic and electrophoretic forces are indicated (EOF is dominant here), middle panels present characteristic current blockades caused by analyte detection. Analytes were added to *cis* for A2V1 under negative voltage and to *trans* for A6V1 under positive voltage due to its reduced conductance at negative potentials. Right panels: dwell time (ln[t]) vs. voltage plots. A) Detection of polypeptide ‘tzatziki’ (10 µL addition of elution sample). Upon increasing voltages, average dwell times (t) were reduced, confirming translocation. Buffer conditions: 1 M KCl, 15 mM TRIS, pH 7.5. Scale bar: 0.1 s. B) MBPa translocation driven by electroosmotic forces. The reducing dwell time upon increasing voltage demonstrates successful translocation of MBPa proteins. The increase in dwell time with A6V1 at +200 mV shows obstruction in the nanopore. Buffer conditions: 1 M KCl, 50 mM citric acid, 2 M urea and pH 4.5. Scale bar: 0.1 s. All measurements performed in triplicate. Measurements were recorded with 50 kHz sample frequency and 10 kHz low-pass Bessel filter.

Finally, we investigated the possibility to translocate full proteins with A2V1 in PBD_11_PEO_8_ and A6V1 in PBD_22_PEO_14_. Earlier work reported translocation of full-length Gu.HCl-unfolded proteins through α-hemolysin nanopores inserted in PBD_11_PEO_8_ bilayers.^[35]^ The polymer membrane was found to be beneficial due to its stability towards high concentrations of the denaturing agent (guanidinium chloride) required for translocation.^[18]^ Previously, we studied the translocation of urea-unfolded proteins using CytK-4D in DPhPC.^[36]^ We found that maltose binding proteins (MBPa, MBP containing a N_10_LGIEGLYFQSH tail – a spacer polypeptide used for a different study – at the C-terminus, and a His_6_-tag at the N-terminus) in mild unfolding conditions (1 M KCl, 50 mM citric acid, 2 M urea and pH 4.5) could also translocate across the nanopore. Here, MBPa (5 µM, *cis*) events were observed with A2V1 in PBD_11_PEO_8_ above a threshold potential of −140 mV (**Figure 4B**). Under −140 mV, the dwell time averaged 1.54 ± 0.06 ms and *I*_ex_ = 80.6 ± 3.1 %. Upon increasing the applied potential up to −200 mV, *I*_ex_ remained similar (84.9 ± 2.3 %) whereas the dwell time decreased gradually to 0.90 ± 0.12 ms, thus revealing successful translocation of MBPa by A2V1. Similarly, A6V1 reconstituted in PBD_22_PEO_14_ enabled MBPa detection above voltages of +140 mV (**Figure 4B**, 3 µM, *trans*), with decreasing dwell time values with increased applied bias (1.78 ± 0.02 ms at +140 mV and 1.44 ± 0.19 ms at +180 mV), before a substantial dwell time increase occurred to 2.94 ± 0.83 ms under +200 mV. Moreover, *I*_ex_ remained roughly the same (84.0 ± 3.1% at +140 mV, 83.4 ± 3.5% at +160 mV and 83.1 ± 3.42% at +180 mV), only increasing under +200 mV (89.4 ± 1.9%). The increase in dwell time and *I*_ex_ at +200 mV suggests that under these conditions the transport of the unfolded protein may not take place. The obstruction is possibly caused by blob-formation, obstruction of the nanopore by coiled protein domains occurring under conditions of high electro-osmotic flow^[37]^, or by interaction with PEO groups in the polymer membrane. However, despite showing higher translocation potential and lower *I*_ex_, this work shows that the extended nanopores can be used to translocate unstructured proteins.

## Discussion and Conclusions

The incorporation of nanopores in portable biosensing devices requires robust amphiphilic membranes, stable to mechanical perturbations and biological samples. While amphiphilic polymers can provide this required stability, long-term nanopore reconstitution into the synthetic membrane can be cumbersome or not possible given the increased thickness and changed chemical interactions compared to lipid-based native membranes. This is a significant drawback for the development of portable nanopore devices, which require stable insertion to allow long-term use of nanopores, shipment and storage of devices.

To address this shortcoming, we hypothesised that an extension of the transmembrane domain (TMD) in β-barrel nanopores can enhance stable reconstitution in PBD_11_PEO_8_ and PBD_22_PEO_14_ polymer membrane, as it would reduce the hydrophobic mismatch occurring between the TMD and the hydrophobic region of the membrane core. In this work, we engineered new β-barrel nanopores with elongated TMDs, via pairwise addition of pairs of amino acids (AAs) with alternate hydrophobic aromatic side chains. Additional amino acids included glycine residues in the middle of the β-barrel, which are typically found in transmembrane domains of nanopores; and aromatic residues near the region interacting with hydrophilic headgroup of the biological membranes.

CytK constructs easily tolerated the addition of AA pairs. A total of 13 constructs were tested for their electrical properties both in DPhPC and PBD-PEO bilayers, containing 2 extra amino acid per strand in the TMD or ∇2 (+0.7 nm to the TMD, n=2 nanopores), ∇4 (+1.4 nm, n=5), ∇6 (+2.1 nm, n=2), ∇8 (+2.8 nm, n=2) or ∇10 (+3.5 nm, n=2). We found that both ∇2 nanopores inserted well into the PBD_11_PEO_8_ membranes, with stable open pore currents and low noise. By contrast, among the five ∇4 nanopores, only one construct showed favourable current properties (A4V1). This nanopore was the only construct that had an additional glycine residue in the middle of the TMD, and contained a pair-extension on the hydrophilic domain, likely promoting stable interactions with the PEO region on the *cis* side (**Figure 1**). Among the six nanopores designed to fit into in PBD_22_PEO_14_ membranes (∇6, ∇8, and ∇10) only one of the two ∇6 nanopore showed favourable ionic current properties (A6V1). As observed for ∇4 nanopores, the nanopore had an additional glycine residue in the middle of the TMD and extended above the hydrophilic region on the *cis* side. It is surprising that A6V1 nanopore yielded the best results in PBD_22_PEO_14_ bilayers because the ∇6 extension yielded a TMD considerably shorter than the hydrophobic thickness of the membrane.

Molecular dynamics simulations focusing on A2V1 in PBD_11_PEO_8_ and A6V1 in PBD_22_PEO_14_ showed that these constructs require substantial deformation from the polymer membrane for reconstitution. In turn, the resulting membrane curvature exposed the trans entry of the nanopore to PEO-rich regions. The PEO layer was observed to protrude into the nanopore, especially in the A6V1 construct, and to slow down potassium transport through specific interaction. Together with the longer resistance produced by the elongated TMD in the polymer membrane, it provided a likely molecular explanation for the experimentally observed reduction in conductivity compared to nanopores embedded in lipid bilayers.

Nanopore constructs yielding favourable current properties in PBD-PEO bilayers (A2V1, A2V2 and A4V1 in PBD_11_PEO_8_, A6V1 in PBD_22_PEO_14_) showed the ability to transiently bind to γ-cyclodextrins, thus demonstrating similar nanopore functioning as in lipid membranes. Expectedly, subtle changes were observed compared to the original CytK-4D nanopore, possibly related to the increased length of the nanopore and a slightly different tilt of the β-barrel in the membrane. Importantly, the extended nanopores maintained a strong electroosmotic flow, which allowed successful translocation of both unstructured 140 AA model polypeptides and full-length maltose binding proteins in mild denaturing conditions. The stabilisation of these nanopores, especially those with enhanced EOF, in robust polymer membranes is therefore a promising step towards the realisation of new nanopore-based biosensing devices.

## Materials and Methods

### Chemicals and reagents

The chemicals and suppliers used are listed as follow: Amplicillin sodium salt was purchased from Fisher Bio Reagents; chloramphenicol (≥98.0) from Sigma Life Science); urea (≥99.5%), guanidinium chloride (≥99.5%, biochemistry), isopropylthio-β-galactoside (≥99.0%, dioxin-free, animal-free), LB medium, 2xYT medium, NaCl (≥99.5%), HEPES (PUFFERAN® CELLPURE® (≥99.5%), imidazole (≥99%), KCl (≥99.5%), Tris(2-carboxyethyl)phosphine hydrochloride (≥98.0%), Dodecyl-β-D-maltosid (≥99%) from Roth; citric acid (≥99.6%, anhydrous and n-hexadecane (99% from Acros Organics; BIS-TRIS propane (≥99.0%) from Sigma Life Sciences; protease inhibitors (Pierce™ Protease inhibitor Mini tablets, EDTA-free); GeneJET gel extraction kit, GeneJET PCR purification kit, GeneJET Plasmid Miniprep kit were purchased from (Thermo Scientific); Ni-NTA agarose from Qiagen; DPhPC from Avanti polar lipids; PBD-PEO from Polymer Source, Inc.; n-pentane from Sigma-Aldrich; DNA primers and gBlock™ from IDT. BL21(DE3) strain harbouring the pET-PfuX7 plasmid was kindly provided by prof. dr. Oscar Kuipers.

### Cloning barrel extension variants

Two strategies were employed in the preparation of the barrel extension constructs. In the case of the A4V1, A4V4, A8V1 and A10V1 constructs, the extension was obtained by means of fusion PCR of two amplified fragments and a gblock synthetic DNA fragment. PCRs were performed as previously described.^[23]^ In short, the upstream gene fragment (corresponding to the N-terminal part of the protein preceding the barrel) was amplified with T7pro primer and a primer binding upstream of E112, and the downstream gene fragment (corresponding to the C-terminal part of the protein following the barrel) was amplified with a primer binding downstream of K155D and T7term primer. The two fragments were gel extracted using the GeneJet gel extraction kit. The full gene encoding for the modified nanopore was obtained by fusing the two amplification fragments and the gblock (corresponding to the modified barrel) in the presence of U-containing primers. The 10-14 bps homology regions necessary for USER cloning were selected between the T7pro and ATG codon (upstream) and between the stop codon and T7term (downstream). The full gene was introduced in the pT7-sc1 plasmid by USER cloning, followed by transformation into chemically competent E. coli cells, as previously described.^[23,38]^ Transformants were selected on LB-agar plates supplemented with 100 μg/mL carbenicillin and 1 % glucose.

In the case of the other constructs (A2V1, A2V2, A4V2, A4V3, A4V5, A6V1, A6V2, A8V2, A10V2), the barrel extension was obtained by introducing three U-containing DNA fragments into the pT7-sc1 vector directly by means of USER cloning. In this case, the DNA encoding for the new amino acids was introduced as overhangs on primers bearing uracil at 5’. The four homology regions were chosen as follows: the upstream and downstream HRs were the same as above; the other two, midstream 1 and midstream 2, were designated at the insertion sites within the DNA sequence corresponding to the N-terminal and C-terminal β-strand, respectively. The resulting three DNA fragments were clean-up from the PCR mix using the GeneJet PCR clean up kit. Finally, the plasmids bearing the extension constructs were prepared as above.

### Nanopore and substrate production

The nanopores and substrates were prepared in the same manner as the original CytK-4D nanopore, as described in previous work.^[23]^ Concentration of the tzatziki polypeptide could not be determined via Bradford assays, thus the added volume of elution sample is indicated. For MBP translocation, a slightly different version than the wild-type was used (MBPa), containing a N_10_LGIEGLYFQSH tail at the c-terminus and a His_6_-tag at the N-terminus.

### Membrane formation

Experiments were performed in a custom-built experimental device, comprising two adjacent chambers (max. volume 700 µL each) separated by a 25-µm thick Teflon sheet containing an aperture (100‒150 µm in diameter). Vertically free-standing membranes were formed across this aperture using the Montal-Mueller technique.^[39,40]^ The same protocol was followed for bilayer formation with PBD-PEO polymers as for DPhPC lipids, described in detail in earlier work.^[19]^

### Data acquisition and analysis

Electrophysiology was performed with a setup comprising an Axopatch 200B patch clamp amplifier and DigiData 1440 A/D converter, using Clampex 10.7 software provided by Molecular Devices. Measurements for current comparisons were taken using 10 kHz sample frequency and 2 kHz Bessel low-pass filter. Open pore current values (*I*_0_) of each nanopore construct were average values taken from 3 individually inserted nanopores. *I*_0_ was obtained from a Gaussian fit taken on a histogram (bin size 0.5 pA) of all individual current values measured over 10 seconds (−75 mV for constructs A2V1‒A4V5, +75 mv for constructs A6V1‒A8V2, buffer: 1 M KCl, 15 mM TRIS, pH 7.5). All single-molecule experiments were carried out using 50 kHz sampling frequency and 10 kHz Bessel low-pass filter. Excluded current (*I*_ex_) calculations were calculated as follows: 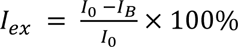, with *I*_B_ being the blocked current value during an event. For event detection, events were included that induced a current blockade exceeding a threshold value of 5× the standard error of the open pore current. To determine the average dwell time for γ-CD and MBP measurements, values from events were grouped in conventional histograms and fitted with an exponential fit to obtain the average dwell time (bin width at least 3× smaller than resulting average dwell time). For tzatziki measurements, events were grouped in an exponential histogram and fitted with a Gaussian fit to obtain log[dwell time] (value for bin size was iteratively optimised until the histogram resembled a Gaussian distribution). Average *I*_ex_ values were obtained by grouping data in a conventional histogram and fitting with a Gaussian fit (bin size 0.5 pA). All measurements were performed in (at least) triplicates. The minimum amount of data points acquired with tzatziki measurements at the lowest applied potential were as follows: 415 (A2V1, −120 mV), 827 (A2V2, −140 mV), 916 (A4V1, −140 mV), all in PBD_11_PEO_8_ and 126 (A6V1,in PBD_22_PEO_14_, +140 mV). Because of the limited data points acquired per individual measurement for A6V1 constructs, a total of 6 individual measurements were included, yielding on average 290 events per measurement. MBP measurements with A2V1 contained at least 251 events at the lowest potential (−140 mV) while for A6V1 this included 260 events (+140 mV). γ-cyclodextrin measurements included the following minimum number of events per construct: A2V1 1678; A2V2 1649; A4V1 957 (all in PBD_11_PEO_8_); A6V1 329 (in PBD_22_PEO_14_).

### Molecular Dynamics Simulations

The initial protein structure of the A2V1 pore was obtained using the SwissModel web server.^[41]^ The Cyt4K homology model, which accompanied the crystal structure, was used as a template. The A6v1 nanopore coordinates were obtained using Alphafold2.^[42]^ All molecular dynamics simulations were performed using GROMACS (version 2023.4).^[43]^ Simulations at the all-atom level utilized the CHARMM36m force field (version July2022) for proteins and ions, as well as CHARMM TIP3P water.^[44,45]^ CHARMM parameters for DPhPC were published in the literature.^[46]^ PEO-b-PBD parameters were taken from a previous study.^[19,47]^ All systems were initially built at the coarse-grained Martini level^[48]^ using TS2CG^[49]^ and a salt concentration of 150mM. Protein parameters were obtained using martinize2^[50]^. The water interaction of the protein backbone beads, whose secondary structure was classified as beta-strands, was reduced by 1 kJ/mol to ensure stable insertion of the pore. Water interaction of the backbone beads classified as coil was increased by 0.5 kJ/mol. These adjustments follow the principles of secondary-structure-specific rescaling used to address some shortcomings of the Martini protein model.^[51]^ Subsequently, the systems were relaxed and then equilibrated for 1 µs before backmapping to target resolution. Backmapping of the membrane and water was done using polyply^[47]^, while the protein was backmapped using ezAlign.^[52]^ After backmapping, simulations were run for 20ns at the original salt concentration, followed by the addition of salt to reach the target 1M KCl concentration. Another equilibration cycle was done before a production run was launched.

All equilibrations and production runs were performed under constant temperature at 298.15 K using the v-rescale temperature coupling (τ = 1 ps) with a coupling group for solvent, protein, and membrane.^[53]^ The pressure was kept constant at 1 bar using the Parrinello–Rahman semi-isotropic pressure coupling algorithm (τ = 5 ps, β = 4.5 × 10–5 bar–1) in production and the Berendsen pressure coupling during equilibration.^[54,55]^ Simulation lengths, system sizes, and compositions are listed in Table S4. The MD data was analyzed with the help of MDAnlaysis^[56]^, FreudAnalysis^[57]^, and custom scripts.

The ion mobility profile as a function of z-coordinate and distance from the pore centre was computed as follows: First, trajectories were centred on the protein. Then, for every frame, the potassium within a cylindrical selection of 10nm from the centre of the pore was replaced by a Gaussian density, and the intensity was collected on a 3D grid using FreudAnalysis.

The cantre of the pore was computed by finding the center of the following residues for the A2V1 and A6V1 pores, respectively: THR163, THR115. The cylinder was 1.5 nm wide in both cases. Each voxel in the grid has a size of about 1 potassium ion. From this data, an autocorrelation time as a function of lag time was computed for each voxel using the Fourier transform implemented in *scipy* signal. The obtained autocorrelation time curve was fitted to an analytical autocorrelation function taken from single-molecule fluorescence spectroscopy^[58]^:

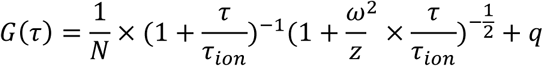

The autocorrelation function does not contain a flow term because the EOF of ions is orders of magnitude smaller than the diffusive component of the motion. This timescale difference makes it unreliable to separate the two components. Furthermore, we include the *q* fudge factor to account for cases where the intensity does not fully decay to zero. In general, it is very small and only applies to a few voxels. Fitting this autocorrelation function yields τ_*ion*_, which is a direct measure of the ion mobility at each voxel. The mobility data is finally averaged radially from the pore centre axis and normalized by the τ_*ion*_computed in the bulk water phase. Thus, it measures the local ion mobility relative to the bulk and allows to identify regions of slow transport. The statistical error was computed by splitting the trajectory into five blocks and averaging the τ_*ion*_values.

## Acknowledgements

The experimental part was financially supported by NWO VICI (no. 192068).

F.G. acknowledges funding from the Klaus Tschira Stiftung gGmbH (Independent PostDoc Fellowship)

## Author contributions

E.V. and A.S. contributed to this work equally and are co-first authors. E.V., A.S. and G.M. designed the experiments. Nanopore constructs were expressed, purified and oligomerised by A.S.; E.V. performed electrical characterisation of the nanopore constructs and single-molecule assays. Data analysis from experiments were performed by both A.S. and E.V.. F.G. designed, performed and analysed the molecular dynamics simulations, A.H. supported F.G. during this work. E.V., A.S., F.G. and G.M. contributed to the writing of the manuscript.

## Conflict of interest

Giovanni Maglia is a founder, director, and shareholder of Portal Biotech Limited, a company engaged in the development of nanopore technologies. This work was not supported by Portal Biotech Limited.

