## Supplementary Information for "β-barrel nanopores designed for insertion into thick block copolymer membranes"

Figure S12: Translocation of tzatziki across the A2V1 construct from the *cis* side (in $PBD_{11}PEO_8$). 19

Figure S13: Translocation of tzatziki across the A2V2 construct from the *cis* side (in $PBD_{11}PEO_8$). 20

Figure S14: Translocation of tzatziki across the A4V1 construct from the *cis* side (in $PBD_{11}PEO_8$). 21

Figure S15: Translocation of tzatziki across the A6V1 construct from the *trans* side (in $PBD_{22}PEO_{14}$ ........................................ 22

**Table S1.** Amino acid sequences of commonly used β-barrel nanopores

| Nanopore | Sequence |
|---|---|
| CytK (WT) | ETTVTSSVSYQLGGSI**K**ASVTPSGPSGESGATG**Q**VTWSD**S**VSY**K** |
| CytK (4D) | ETTVTSSVSYQLGGSI**D**ASVTPSGPSGESGATG**D**VTWSD**D**VSY**D** |
| α-Hemolysin (WT) | EYMSTLTYGFNGNVTGDDTGKIGGLIGANVSIGHTLK |
| Aerolysin (WT) | YDVTLRYDTATNWSKTNTYGLSEKVTTKNKFKWPLVGETELSIEIAANQSWASQNG GSTTTSLSQSVRP |
| Protective antigen (WT) | DQSTQNTDSQTRTISKNTSTSRTHTSEVHGNAEVHASFFDIGGSVSAGFSNSNSST VAIDHSLSLAGERTWAETMG |
| Lysenin (WT) | VDQKITITKGMKNVNSETRTVTATHSIGSTISTGDAFEIGSVEVSYSHSHEESQVSMTE[illegible] EVYESKVIEHTITI |

blue = lumen-exposed residues
red = *trans* loop

**Table S2.** Amino acid sequences for all designed nanopore constructs using CytK-4D as a template

| Construct | Sequence |
|---|---|
| A2V1 | MAQTTSQVVTDIGQNAKTHTSYNTFNNEQADNMTMSLKVTFIDDPSADKQIAVINT<br>TGSF<br>MKANPTLSDAPVDGYPIPGASVTLRYPSQYDIAMNLQDNTSRFFHVAPTNAVEETT<br>VTSS<br>VSYTYQLGGSIDASVTPSGPSGESGATGDVSVTWSDDVSYDQTSYKTNLIDQTNK<br>HVKWN<br>VFFNGYNNQNWGIYTRDSYHALYGNQLFMYSRTYPHETDARGNLVPMNDLPTLT<br>NSGFSP<br>GMIAVVISEKDTEQSSIQVAYTKHADDYTLRPGFTFGTGNWVGNNIKDVDQKTFNK<br>SFVL<br>DWKNKKLVEKKGSAHHHHHH |
| A2V2 | MAQTTSQVVTDIGQNAKTHTSYNTFNNEQADNMTMSLKVTFIDDPSADKQIAVINT<br>TGSF<br>MKANPTLSDAPVDGYPIPGASVTLRYPSQYDIAMNLQDNTSRFFHVAPTNAVEETT<br>VTSS<br>VSVTYQLGGSIDASVTPSGPSGESGATGDVSYTWSDDVSYDQTSYKTNLIDQTNK<br>HVKWN<br>VFFNGYNNQNWGIYTRDSYHALYGNQLFMYSRTYPHETDARGNLVPMNDLPTLT<br>NSGFSP<br>GMIAVVISEKDTEQSSIQVAYTKHADDYTLRPGFTFGTGNWVGNNIKDVDQKTFNK<br>SFVL<br>DWKNKKLVEKKGSAHHHHHH |
| A4V1 | MAQTTSQVVTDIGQNAKTHTSYNTFNNEQADNMTMSLKVTFIDDPSADKQIAVINT<br>TGSF<br>MKANPTLSDAPVDGYPIPGASVTLRYPSQYDIAMNLQDNTSRFFHVAPTNAVEETT<br>VTSS<br>VTVSYQLGLGGSIDASVTPSGPSGESGATGTGDVTWSVSDDVSYDQTSYKTNLID<br>QTNKH<br>VKWNVFFNGYNNQNWGIYTRDSYHALYGNQLFMYSRTYPHETDARGNLVPMND<br>LPTLTNS<br>GFSPGMIAVVISEKDTEQSSIQVAYTKHADDYTLRPGFTFGTGNWVGNNIKDVDQ<br>KTFNK<br>SFVLDWKNKKLVEKKGSAHHHHHH |
| A4V2 | MAQTTSQVVTDIGQNAKTHTSYNTFNNEQADNMTMSLKVTFIDDPSADKQIAVINT<br>TGSF<br>MKANPTLSDAPVDGYPIPGASVTLRYPSQYDIAMNLQDNTSRFFHVAPTNAVEETT<br>VTSS<br>VSYTVSYQLGGSIDASVTPSGPSGESGATGDVTYSVTWSDDVSYDQTSYKTNLID<br>QTNKH<br>VKWNVFFNGYNNQNWGIYTRDSYHALYGNQLFMYSRTYPHETDARGNLVPMND<br>LPTLTNS<br>GFSPGMIAVVISEKDTEQSSIQVAYTKHADDYTLRPGFTFGTGNWVGNNIKDVDQ<br>KTFNK<br>SFVLDWKNKKLVEKKGSAHHHHHH |
| A4V3 | MAQTTSQVVTDIGQNAKTHTSYNTFNNEQADNMTMSLKVTFIDDPSADKQIAVINT<br>TGSF |

| | |
|---|---|
| | MKANPTLSDAPVDGYPIPGASVTLRYPSQYDIAMNLQDNTSRFFHVAPTNAVEETT<br>VTSS<br>VSVTYSYQLGGSIDASVTPSGPSGESGATGDVTVSYTWSDDVSYDQTSYKTNLID<br>QTNKH<br>VKWNVFFNGYNNQNWGIYTRDSYHALYGNQLFMYSRTYPHETDARGNLVPMND<br>LPTLTNS<br>GFSPGMIAVVISEKDTEQSSIQVAYTKHADDYTLRPGFTFGTGNWVGNNIKDVDQ<br>KTFNK<br>SFVLDWKNKKLVEKKGSAHHHHHH |
| A4V4 | MAQTTSQVVTDIGQNAKTHTSYNTFNNEQADNMTMSLKVTFIDDPSADKQIAVINT<br>TGSF<br>MKANPTLSDAPVDGYPIPGASVTLRYPSQYDIAMNLQDNTSRFFHVAPTNAVEEV<br>STTVT<br>SSVSYQLGGSISIDASVTPSGPSGESTSGATGDVTWSDDVSYTVDQTSYKTNLID<br>QTNKH<br>VKWNVFFNGYNNQNWGIYTRDSYHALYGNQLFMYSRTYPHETDARGNLVPMND<br>LPTLTNS<br>GFSPGMIAVVISEKDTEQSSIQVAYTKHADDYTLRPGFTFGTGNWVGNNIKDVDQ<br>KTFNK<br>SFVLDWKNKKLVEKKGSAHHHHHH |
| A4V5 | MAQTTSQVVTDIGQNAKTHTSYNTFNNEQADNMTMSLKVTFIDDPSADKQIAVINT<br>TGSF<br>MKANPTLSDAPVDGYPIPGASVTLRYPSQYDIAMNLQDNTSRFFHVAPTNAVEETT<br>VTSS<br>VTVSYGIQLGGSIDASVTPSGPSGESGATGDGTVTVSWSDDVSYDQTSYKTNLID<br>QTNKH<br>VKWNVFFNGYNNQNWGIYTRDSYHALYGNQLFMYSRTYPHETDARGNLVPMND<br>LPTLTNS<br>GFSPGMIAVVISEKDTEQSSIQVAYTKHADDYTLRPGFTFGTGNWVGNNIKDVDQ<br>KTFNK<br>SFVLDWKNKKLVEKKGSAHHHHHH |
| A6V1 | MAQTTSQVVTDIGQNAKTHTSYNTFNNEQADNMTMSLKVTFIDDPSADKQIAVINT<br>TGSF<br>MKANPTLSDAPVDGYPIPGASVTLRYPSQYDIAMNLQDNTSRFFHVAPTNAVEETT<br>VTSS<br>VDVSYTYQLGLGGSIDASVTPSGPSGESGATGTGDVSVTWSVSDDVSYDQTSYK<br>TNLIDQ<br>TNKHVKWNVFFNGYNNQNWGIYTRDSYHALYGNQLFMYSRTYPHETDARGNLVP<br>MNDLPT<br>LTNSGFSPGMIAVVISEKDTEQSSIQVAYTKHADDYTLRPGFTFGTGNWVGNNIKD<br>VDQK<br>TFNKSFVLDWKNKKLVEKKGSAHHHHHH |

Continued on next page

| | |
|---|---|
| A6V2 | MAQTTSQVVTDIGQNAKTHTSYNTFNNEQADNMTMSLKVTFIDDPSADKQIAVINT<br>TGSF |

| | |
|---|---|
| | MKANPTLSDAPVDGYPIPGASVTLRYPSQYDIAMNLQDNTSRFFHVAPTNAVEETT<br>VTSS<br>VSVDYSGTYQLGGSIDASVTPSGPSGESGATGDVSATVSYTWSDDVSYDQTSYK<br>TNLIDQ<br>TNKHVKWNVFFNGYNNQNWGIYTRDSYHALYGNQLFMYSRTYPHETDARGNLVP<br>MNDLPT<br>LTNSGFSPGMIAVVISEKDTEQSSIQVAYTKHADDYTLRPGFTFGTGNWVGNNIKD<br>VDQK<br>TFNKSFVLDWKNKKLVEKKGSAHHHHHH |
| A8V1 | MAQTTSQVVTDIGQNAKTHTSYNTFNNEQADNMTMSLKVTFIDDPSADKQIAVINT<br>TGSF<br>MKANPTLSDAPVDGYPIPGASVTLRYPSQYDIAMNLQDNTSRFFHVAPTNAVEETT<br>VTSS<br>VDATVSYQLGASLGGSIDASVTPSGPSGESGATGTATGDVTWSGSVSDDVSYDQ<br>TSYKTN<br>LIDQTNKHVKWNVFFNGYNNQNWGIYTRDSYHALYGNQLFMYSRTYPHETDARG<br>NLVPMN<br>DLPTLTNSGFSPGMIAVVISEKDTEQSSIQVAYTKHADDYTLRPGFTFGTGNWVGN<br>NIKD<br>VDQKTFNKSFVLDWKNKKLVEKKGSAHHHHHH |
| A8V2 | MAQTTSQVVTDIGQNAKTHTSYNTFNNEQADNMTMSLKVTFIDDPSADKQIAVINT<br>TGSF<br>MKANPTLSDAPVDGYPIPGASVTLRYPSQYDIAMNLQDNTSRFFHVAPTNAVEETT<br>VTSS<br>VSVTYDGTISYQLGGSIDASVTPSGPSGESGATGDVTGSATVSYTWSDDVSYDQT<br>SYKTN<br>LIDQTNKHVKWNVFFNGYNNQNWGIYTRDSYHALYGNQLFMYSRTYPHETDARG<br>NLVPMN<br>DLPTLTNSGFSPGMIAVVISEKDTEQSSIQVAYTKHADDYTLRPGFTFGTGNWVGN<br>NIKD<br>VDQKTFNKSFVLDWKNKKLVEKKGSAHHHHHH |
| A10V1 | MAQTTSQVVTDIGQNAKTHTSYNTFNNEQADNMTMSLKVTFIDDPSADKQIAVINT<br>TGSF<br>MKANPTLSDAPVDGYPIPGASVTLRYPSQYDIAMNLQDNTSRFFHVAPTNAVEETT<br>VTSS<br>VDATVSYTYQLGADLGGSIDASVTPSGPSGESGATGTATGDVSVTWSGSVSDDV<br>SYDQTS<br>YKTNLIDQTNKHVKWNVFFNGYNNQNWGIYTRDSYHALYGNQLFMYSRTYPHET<br>DARGNL<br>VPMNDLPTLTNSGFSPGMIAVVISEKDTEQSSIQVAYTKHADDYTLRPGFTFGTGN<br>WVGN<br>NIKDVDQKTFNKSFVLDWKNKKLVEKKGSAHHHHHH |

Continued on next page

| | |
|---|---|
| A10V2 | MAQTTSQVVTDIGQNAKTHTSYNTFNNEQADNMTMSLKVTFIDDPSADKQIAVINT<br>TGSF |

| | |
|---|---|
| | MKANPTLSDAPVDGYPIPGASVTLRYPSQYDIAMNLQDNTSRFFHVAPTNAVEETT<br>VTSS<br>VSVDYSGTISATYQLGGSIDASVTPSGPSGESGATGDVSATGDATVSYTWSDDVS<br>YDQTS<br>YKTNLIDQTNKHVKWNVFFNGYNNQNWGIYTRDSYHALYGNQLFMYSRTYPHET<br>DARGNL<br>VPMNDLPTLTNSGFSPGMIAVVISEKDTEQSSIQVAYTKHADDYTLRPGFTFGTGN<br>WVGN<br>NIKDVDQKTFNKSFVLDWKNKKLVEKKGSAHHHHHH |

**Table S3.** Potassium Current from MD simulations (pA)

| Construct | Membrane | $<I_0>$ ( ~ -400mV) pA | $<I_0>$ ( ~400mV) pA | $R_{sym}$ | $<I_0>$ (~ 200mV) pA | $<I_0>$ (~200 mV) pA | $R_{sym}$ |
|---|---|---|---|---|---|---|---|
| A2V1 | DPhPC | - 859 ± 16 | 1030 ± 27 | 0.84 | -524 ± 12 | 554 ± 12 | 1.05 |
| A2V1 | $PBD_{11}PEO_8$ | -1009 ± 39 | 984 ± 45 | 0.98 | -538 ± 34 | 517 ± 11 | 0.96 |
| A6V1 | DPhPC | -1115 ± 16 | n/a | n/a | -602 ± 21 | n / a | n/a |
| A6V1 | $PBD_{22}PEO_{14}$ | -131 ± 7 | 562 ± 17 | 4.3 | -36 ± 4 | 80 ± 6 | 2.2 |

* The field in MD is box size dependent due to small fluctuations; the fields between the rows are slightly different per row. From top to bottom 406mV, 203mV, 428mV, 214mV, 468mV, 234mV, 426mV, 213mV

**Table S4.** Overview of MD simulations

| Construct | Membrane | Field (V/nm)* | z (nm) | x,y (nm) | run-time (ns) |
|---|---|---|---|---|---|
| A2V1 | DPhPC | 0.02 | 20.01 ± 0.07 | 13.4 ± 0.02 | 164.84 |
| A2V1 | DPhPC | 0.01 | 20.02 ± 0.06 | 13.37 ± 0.02 | 539.52 |
| A2V1 | DPhPC | -0.02 | 20.40 ± 0.04 | 13.33 ± 0.02 | 473.10 |
| A2V1 | DPhPC | -0.01 | 20.3 ± 0.2 | 13.35 ± 0.04 | 510.17 |
| A2V1 | $PBD_{11}PEO_8$ | 0.02 | 21.16 ± 0.09 | 13.00 ± 0.03 | 250 |
| A2V1 | $PBD_{11}PEO_8$ | 0.01 | 21.85 ± 0.07 | 12.84 ± 0.02 | 250 |
| A2V1 | $PBD_{11}PEO_8$ | -0.02 | 21.42 ± 0.06 | 12.91 ± 0.02 | 250 |
| A2V1 | $PBD_{11}PEO_8$ | -0.01 | 21.25 ± 0.07 | 12.96 ± 0.09 | 250 |
| A6V1 | DPhPC | -0.02 | 24.01 ± 0.02 | 13.62 ± 0.02 | 236.12 |
| A6V1 | DPhPC | -0.01 | 24.00 ± 0.1 | 13.63 ± 0.03 | 248.68 |
| A6V1 | $PBD_{22}PEO_{14}$ | 0.02 | 21.46 ± 0.02 | 15.00 + 0.01 | 247.3 |
| A6V1 | $PBD_{22}PEO_{14}$ | 0.01 | 21.22 ± 0.06 | 15.03 ± 0.03 | 216.56 |
| A6V1 | $PBD_{22}PEO_{14}$ | -0.02 | 21.24 ± 0.06 | 15.05 ± 0.03 | 263.9 |
| A6V1 | $PBD_{22}PEO_{14}$ | -0.01 | 21.11 ± 0.05 | 15.10 ± 0.02 | 246.88 |

** the actual field set in the GROMACS mdp settings*

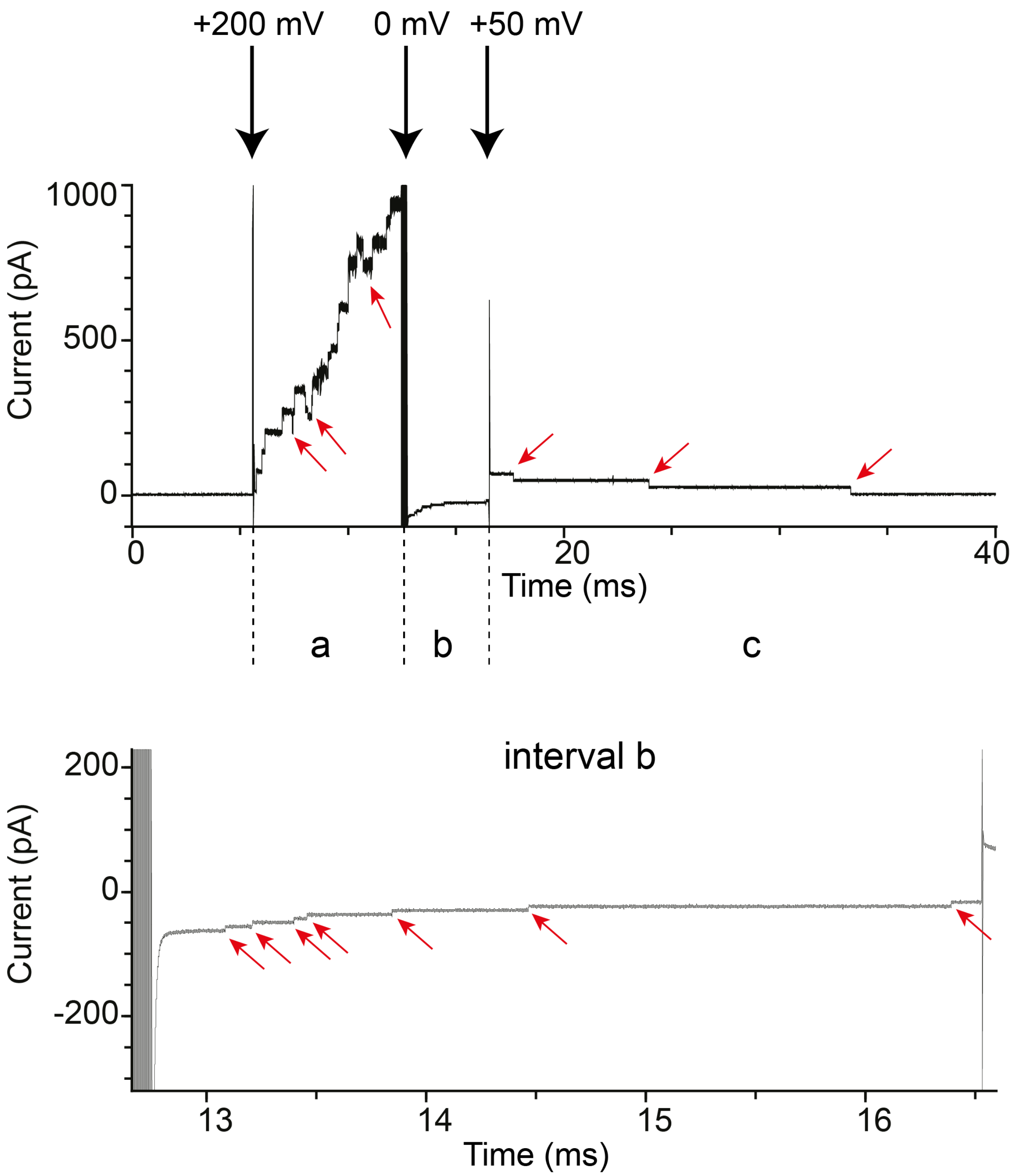

**Figure S1: Insertion of CytK-4D into $PBD_{11}PEO_8$ membrane.** Top trace: Nanopores insert readily when +200 mV is applied (interval a) and occasionally exit (red arrow the membrane). When no voltage is applied (0 mV, interval c) nanopore exits can be observed. At +50 mV (interval c), the few remaining nanopores exit. Bottom trace: close-up in the b interval. Recordings were performed in 1 M KCl, 15 mM Tris, pH 7.5, 10 kHz sampling rate and 2 kHz Bessel filter.

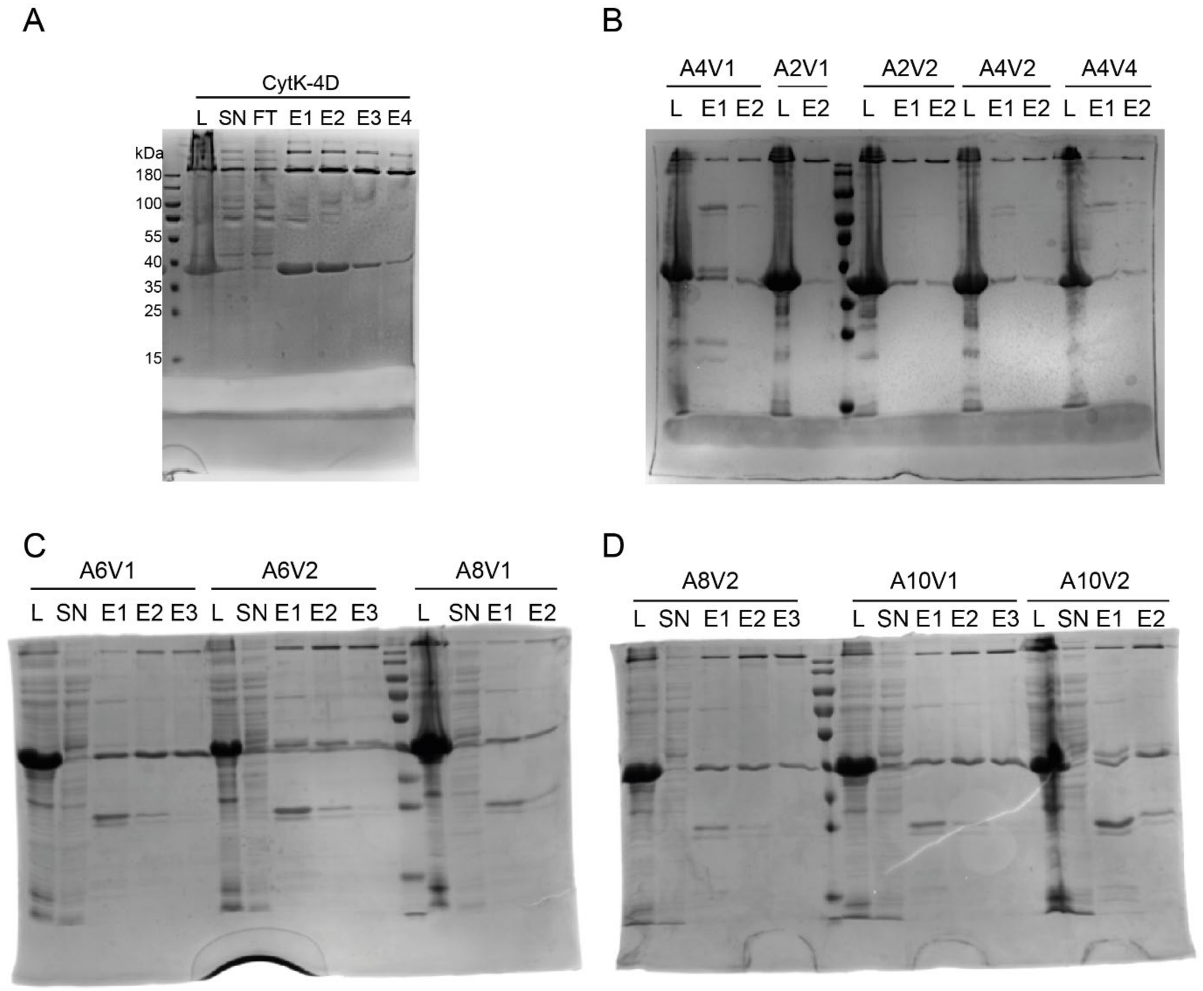

**Figure S2: SDS-PAGE from the purification of the CytK variants.** A) SDS-PAGE of CytK-4D. B-D) SDS-PAGE of the extended constructs. In all cases, both the monomer (expected Mw of 35 to 37 kDa) and the oligomer (expected Mw of 247-259 kDa) can be observed. The expression levels are excellent for all construct. L is the lysate sample, SN is the soluble faction, E1, E2, E3 the elution fractions.

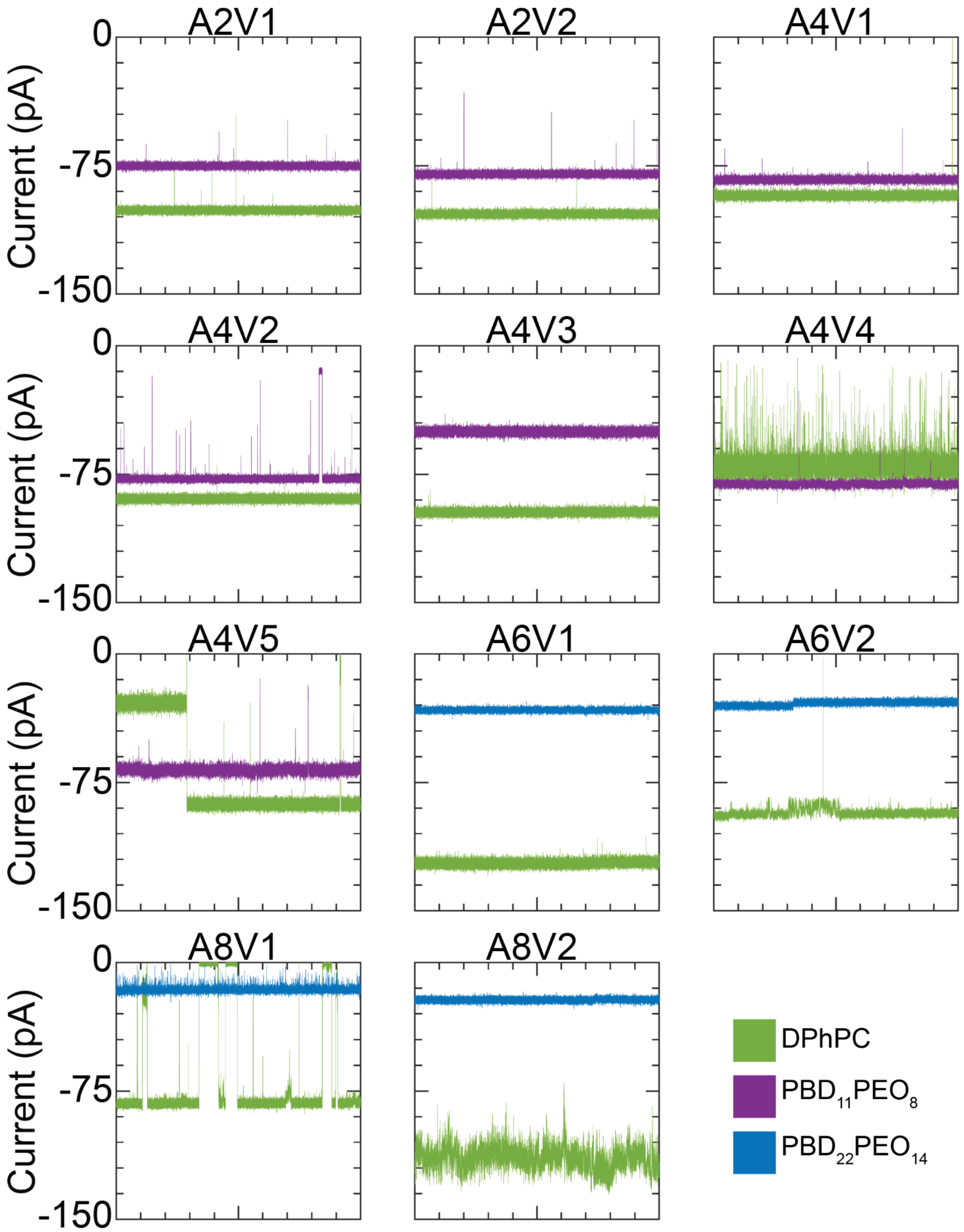

**Figure S3: Representative open pore current ($I_0$) signals of nanopore constructs in DPhPC and PBD-PEO membranes under −75 mV.** Open pore currents are consistently smaller in PBD-PEO membranes. Constructs A2V1–V2, A4V1 and A4V3 show low-noise, stable current signals in $PBD_{11}PEO_8$, although $I_0$ is substantially reduced for A4V3. Constructs A4V4, A6V2 and A8V1–V2 show lower levels of noise and more stable current baselines in $PBD_{11}PEO_8$. A6V1–V2 and A8V1–V2 yield strongly decreased open pore currents under negative voltage in $PBD_{22}PEO_{14}$. All traces show recordings of 10 s. All constructs were tested in triplicate. Buffer conditions: 1 M KCl, 15 mM TRIS, pH 7.5. Measurements were recorded with 10 kHz sample frequency and 2 kHz Bessel low-pass filter.

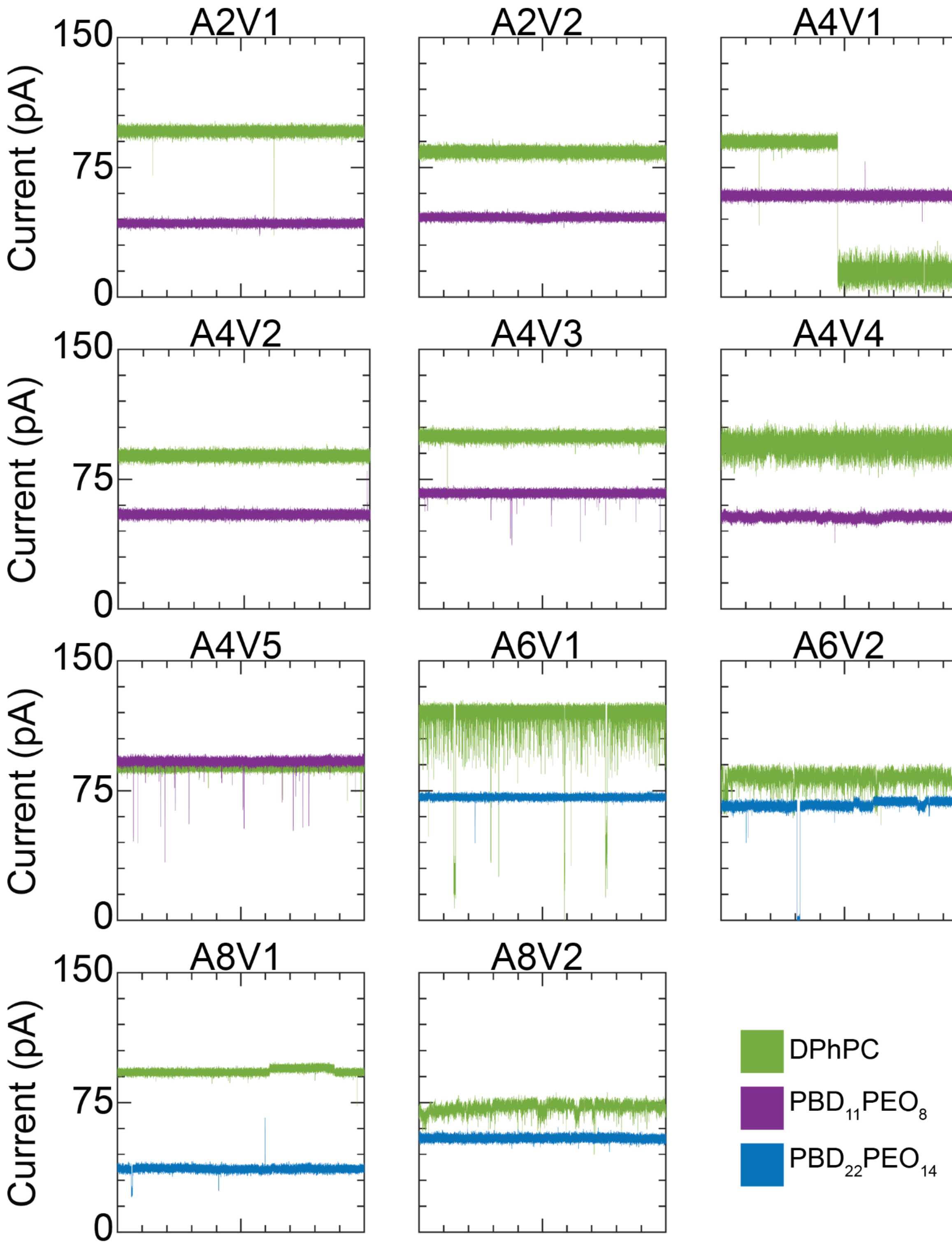

**Figure S4: Representative open pore current ($I_0$) signals of nanopore constructs in DPhPC and PBD-PEO membranes under +75 mV.** $I_0$ values are smaller in PBD-PEO membranes for all constructs except A4V5. Constructs A2V1–V2, A4V1–V3 show low-noise, stable current signals in $PBD_{11}PEO_8$. Constructs A4V4, A6V1–V2 and A8V1–V2 show higher stability in PBD-PEO bilayers, with lower levels of noise and more stable current baselines. A6V1 yields low-noise, stable current signals under positive potential in $PBD_{22}PEO_{14}$. All traces show recordings of 10 s. All constructs were tested in triplicate. Buffer conditions: 1 M KCl, 15 mM TRIS, pH 7.5. Measurements were recorded with 10 kHz sample frequency and 2 kHz Bessel low-pass filter.

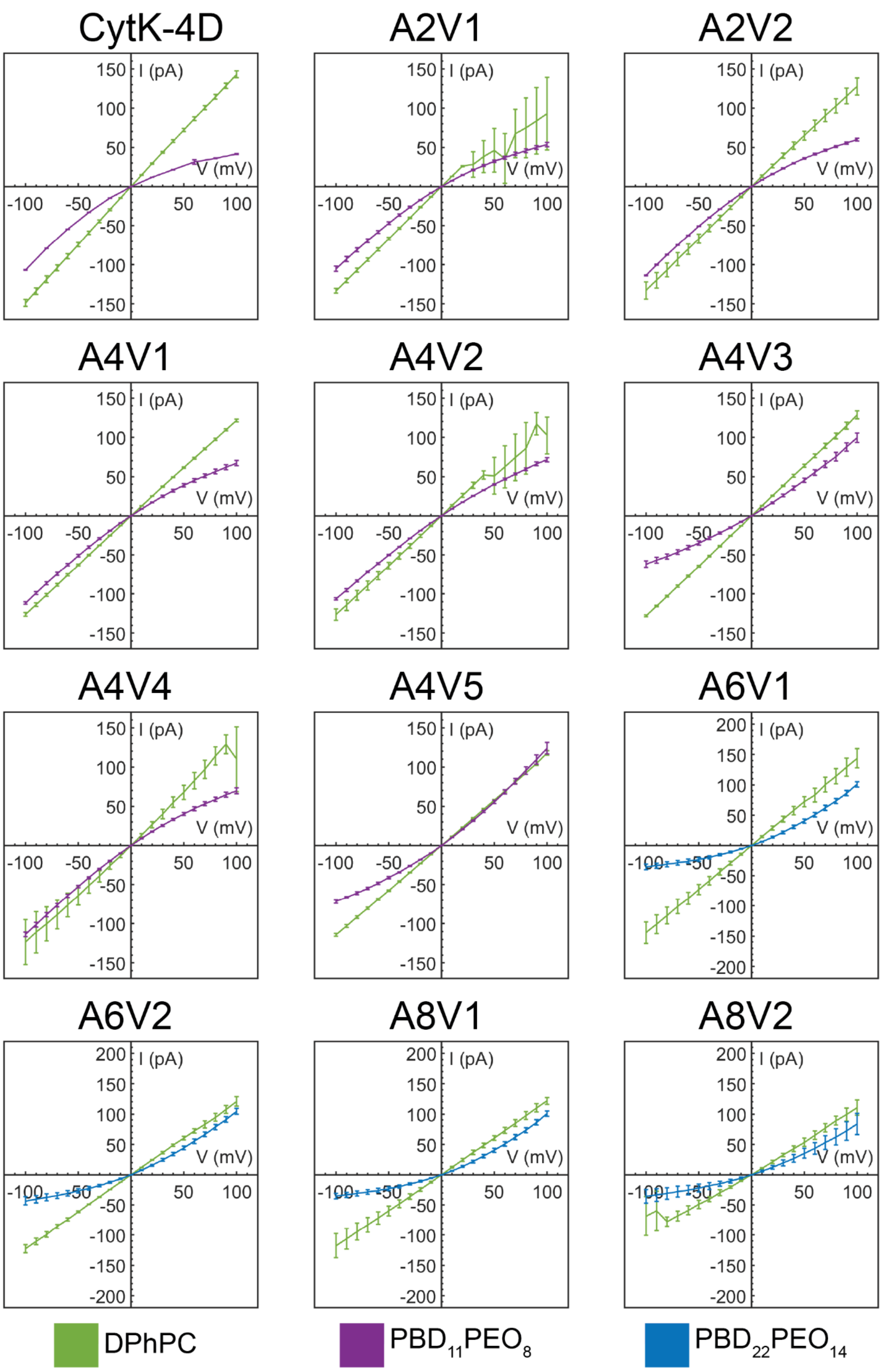

**Figure S5: IV-curves of all characterised nanopore constructs.** IV-curves of CytK-4D and constructs A2V1–A4V5 were measured in DPhPC and PBD$_{11}$PEO$_8$ membranes, constructs A6V1–A8V2 were measured in DPhPC and PBD$_{22}$PEO$_{14}$ membranes. Constructs A2V1–2, A4V1–2 and A4V4 show similar asymmetry in current-voltage dependency as CytK-4D in PBD$_{11}$PEO$_8$, while other constructs show inverted asymmetry. All constructs in PBD$_{22}$PEO$_{14}$ show inverted current-voltage, with low conductance under negative voltages and high conductance under positive voltages. Error bars demonstrate standard deviation (N = 3). Measurements performed in 1 M KCl, 15 mM TRIS, pH 7.5.

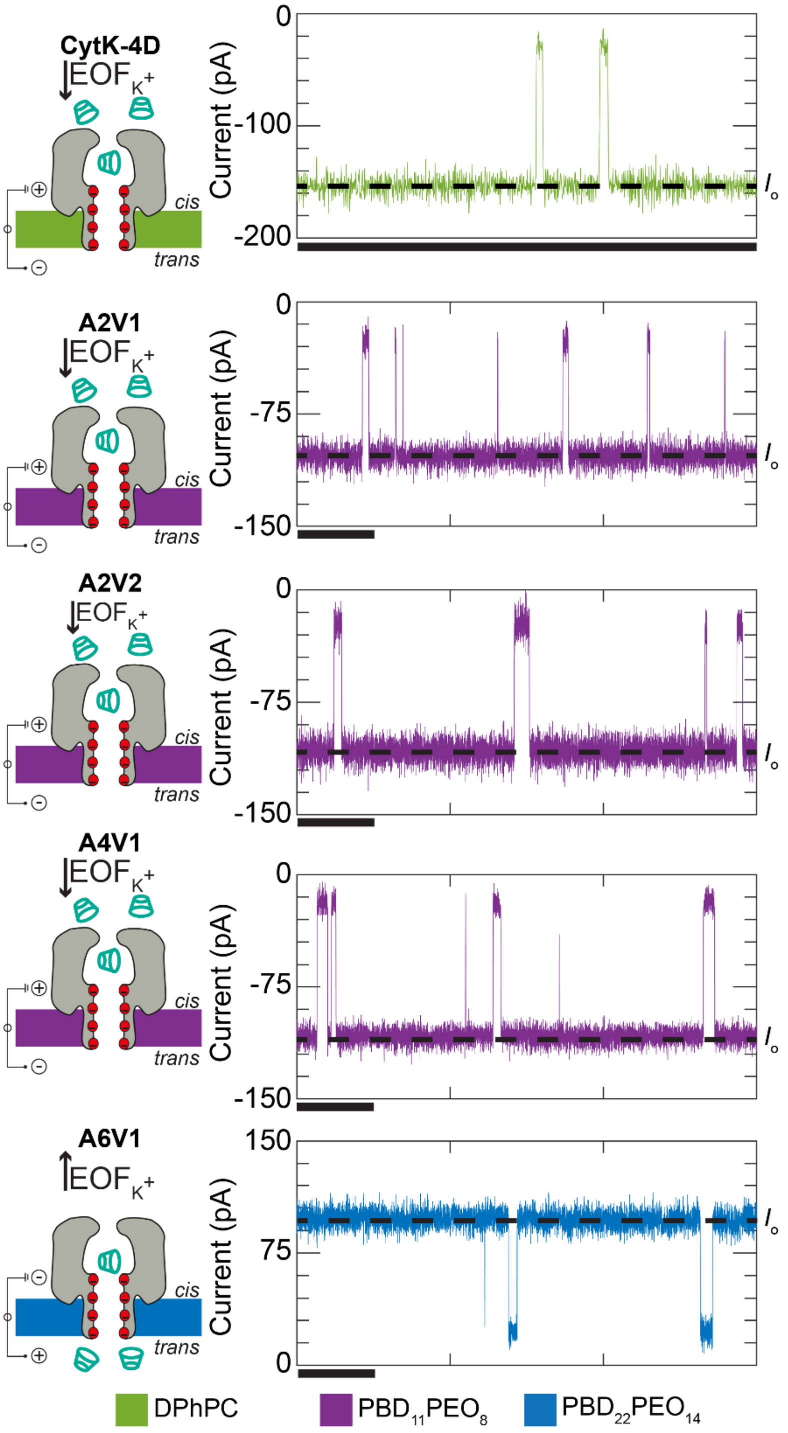

**Figure S6: Detection of γ-cyclodextrin (γ-CD, 5 μM) with CytK-4D and extended nanopore constructs.** Left: schematic of the experimental conditions: CytK-4D in DPhPC, γ-CD added to *cis*; A2V1, A2V2 and A4V1 in $PBD_{11}PEO_8$, γ-CD added to *cis*; A6V1 in $PBD_{22}PEO_{14}$, γ-CD added to *trans*. The direction of the effective electro-osmotic forces (EOF) are indicated with an arrow. Right: individual γ-CD events detected with the nanopore constructs (−100 mV applied to all nanopores except A6V1, +100 mV). Scale bars indicate 25 ms. Buffer contained 1 M KCl, 15 mM TRIS, pH 7.5. Measurements recorded with 50 kHz sample frequency and 10 kHz low-pass Bessel filter. Each measurement was performed in triplicate.

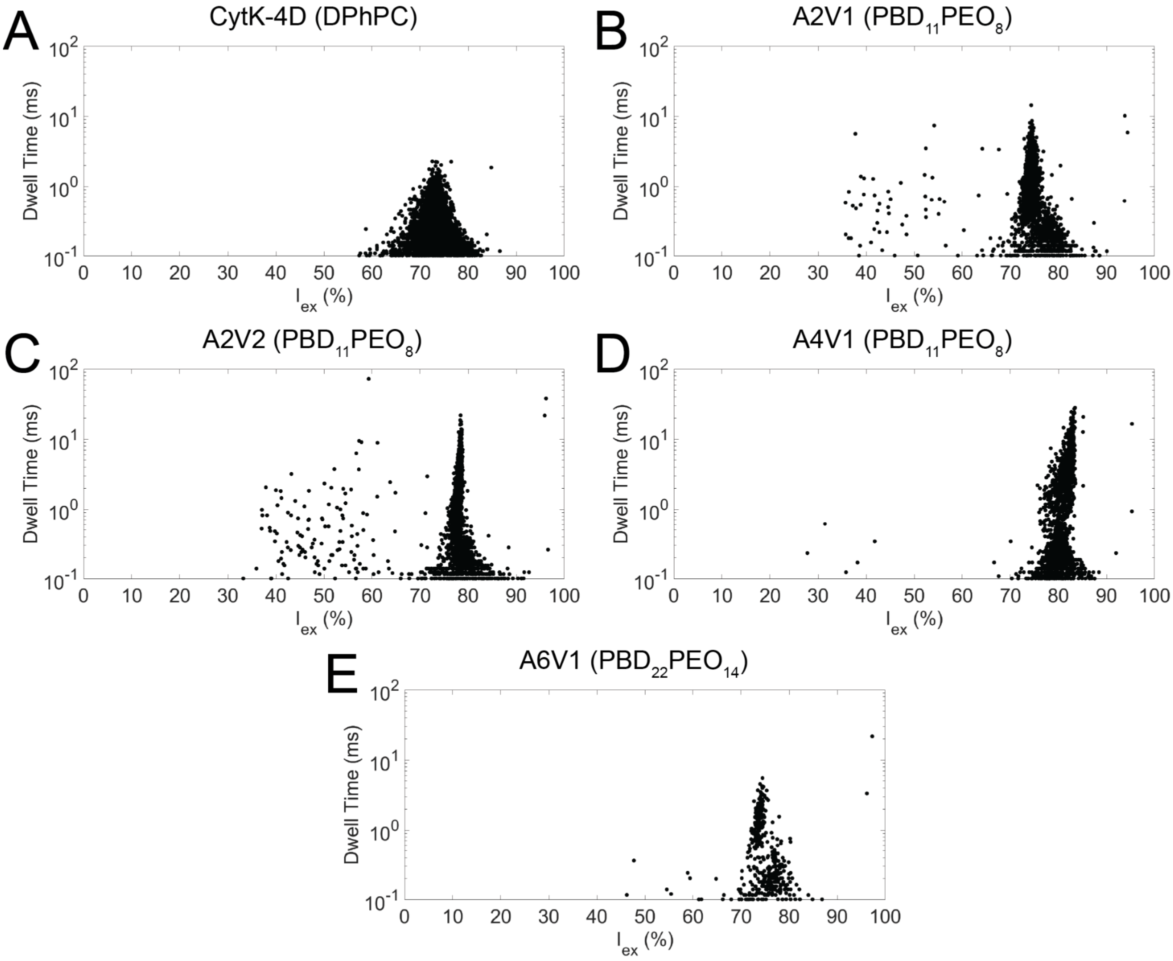

**Figure S7: Representative scatter plots of dwell time versus excluded current (lex) resulting from γ-cyclodextrin (γ-CD, 5 μM) detection with CytK and the extended nanopores in their corresponding membranes.** A) CytK-4D in DPhPC, γ-CD added to *cis* while applying −100 mV. B) A2V1 in $PBD_{11}PEO_8$, γ-CD added to *cis* while applying −100 mV. C) A2V2 in $PBD_{11}PEO_8$, γ-CD added to *cis* while applying −100 mV. D) A4V1 in $PBD_{11}PEO_8$, γ-CD added to *cis* while applying −100 mV. E) A6V1 in $PBD_{22}PEO_{14}$, γ-CD added to *trans* while applying +100 mV. Recording time 60 s. Buffer: 1 M KCl, 15 mM TRIS, pH 7.5. Measurements recorded with 50 kHz sample frequency and 10 kHz low-pass Bessel filter. Measurements were performed in triplicate.

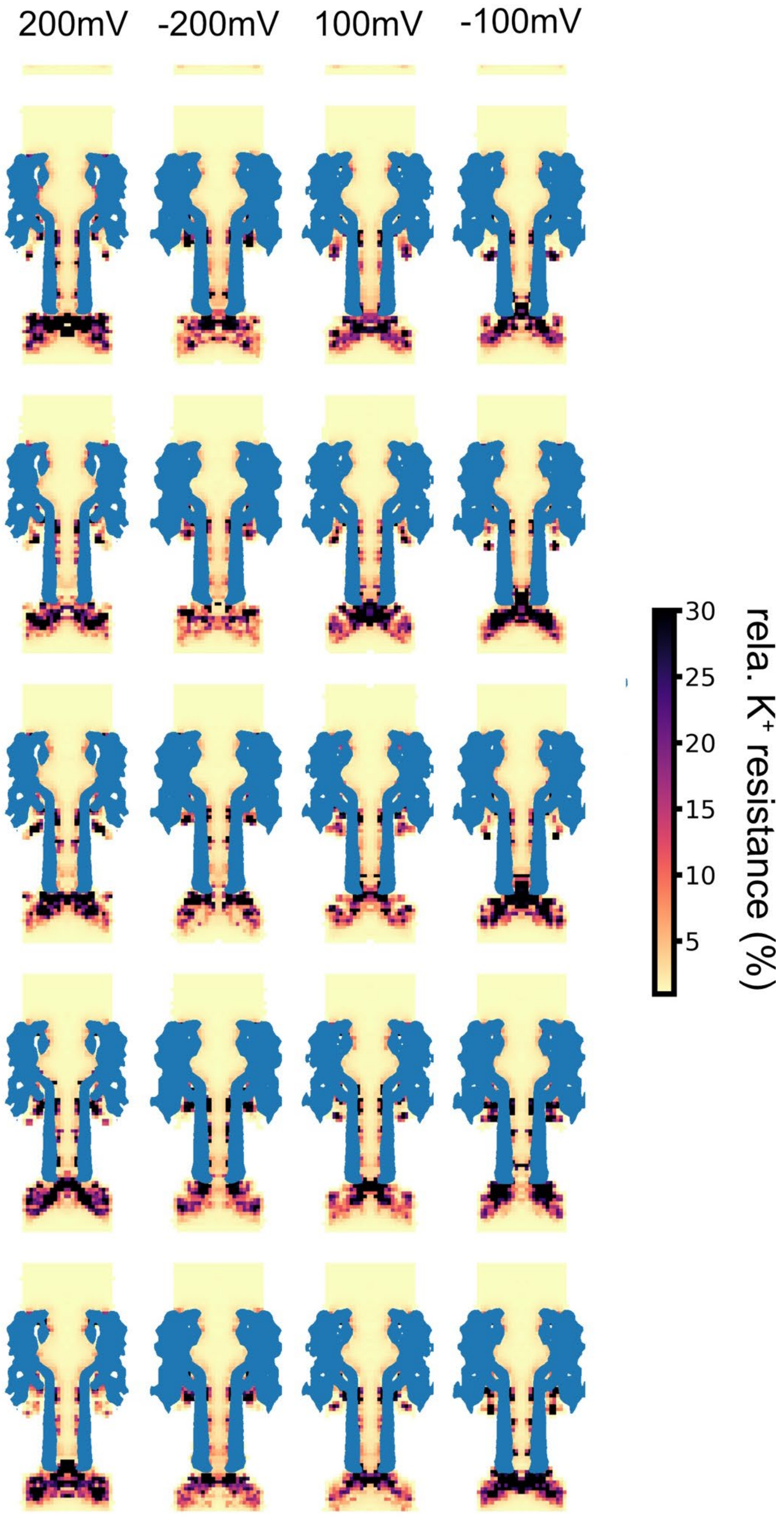

**Figure S8: Relative potassium resistance for the A6V1 construct in $PBD_{22}PEO_{14}$.** Columns show different voltages and rows show blocks of 50ns simulations.

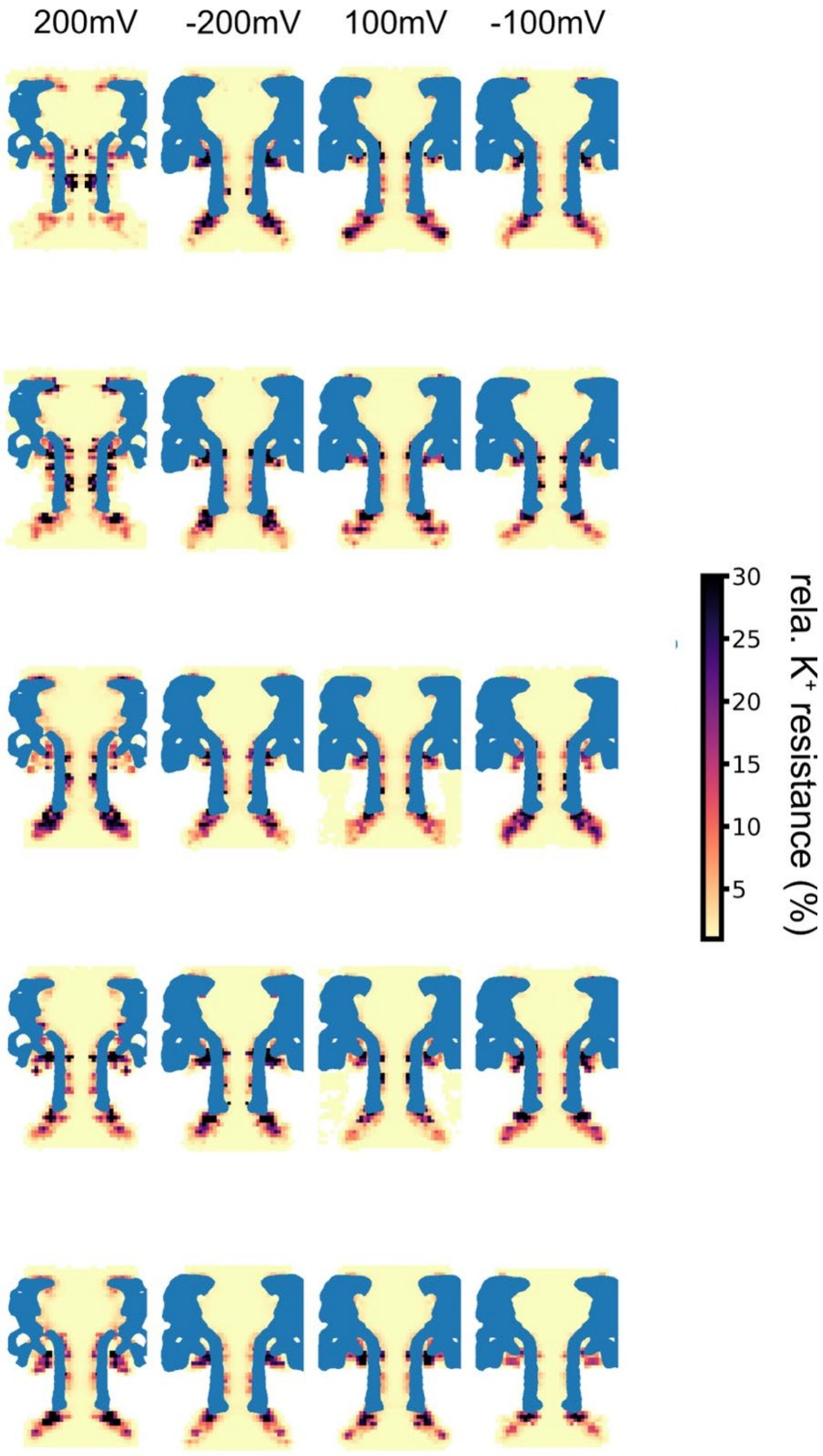

**Figure S9: Relative potassium resistance for the A2V1 construct in $PBD_{11}PEO_8$.** Columns show different voltages and rows show blocks of 50ns simulations.

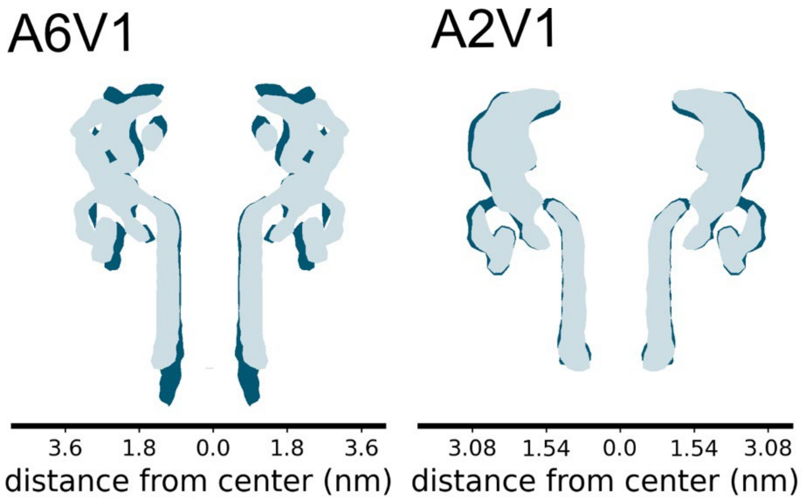

**Figure S10: Backbone density as a function of distance from the centre of the pore for two constructs (A6V1 & A2V1).** The dark blue outline shows the pore in $PBD_{22}PEO_{14}$ and $PBD_{11}PEO_{8}$, respectively, while the light-blue contour corresponds to the pore embedded in a DPhPC lipid bilayer membrane.

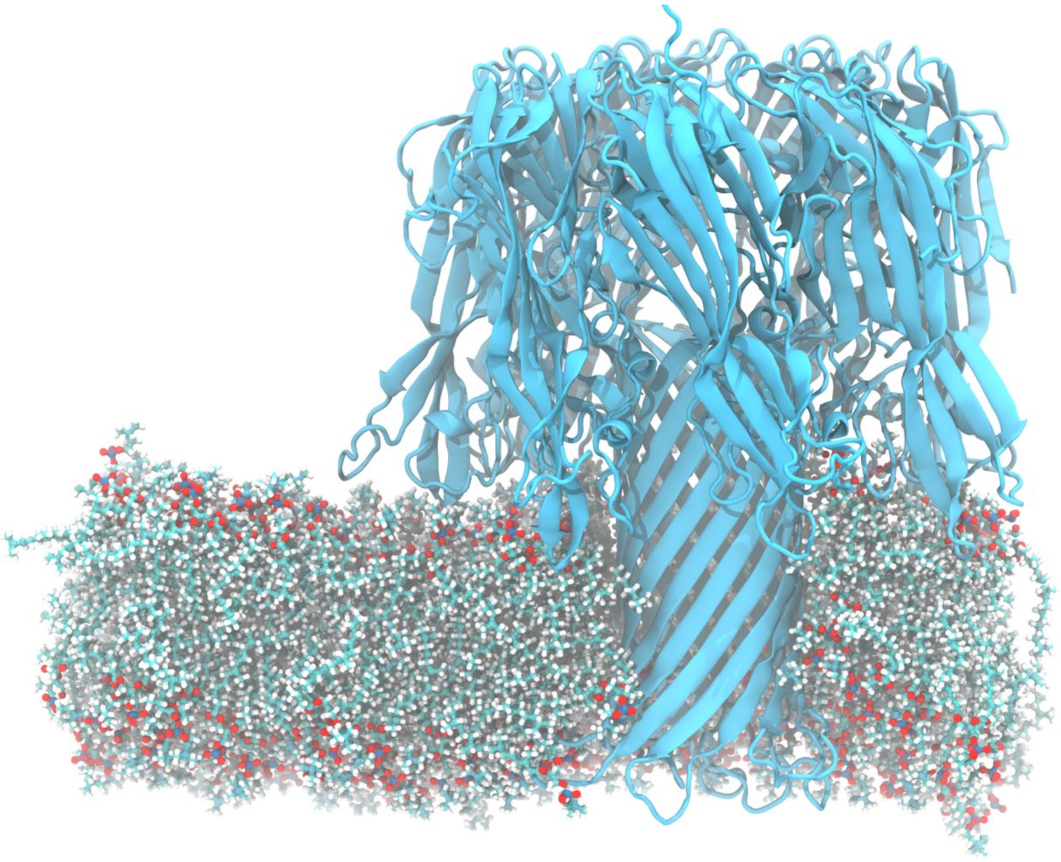

**Figure S11: Snapshot of the last frame of an MD simulation of the A6V1 construct in DPhPC.** The secondary structure is highlighted in cartoon style and was computed using Stride as implemented in VMD.

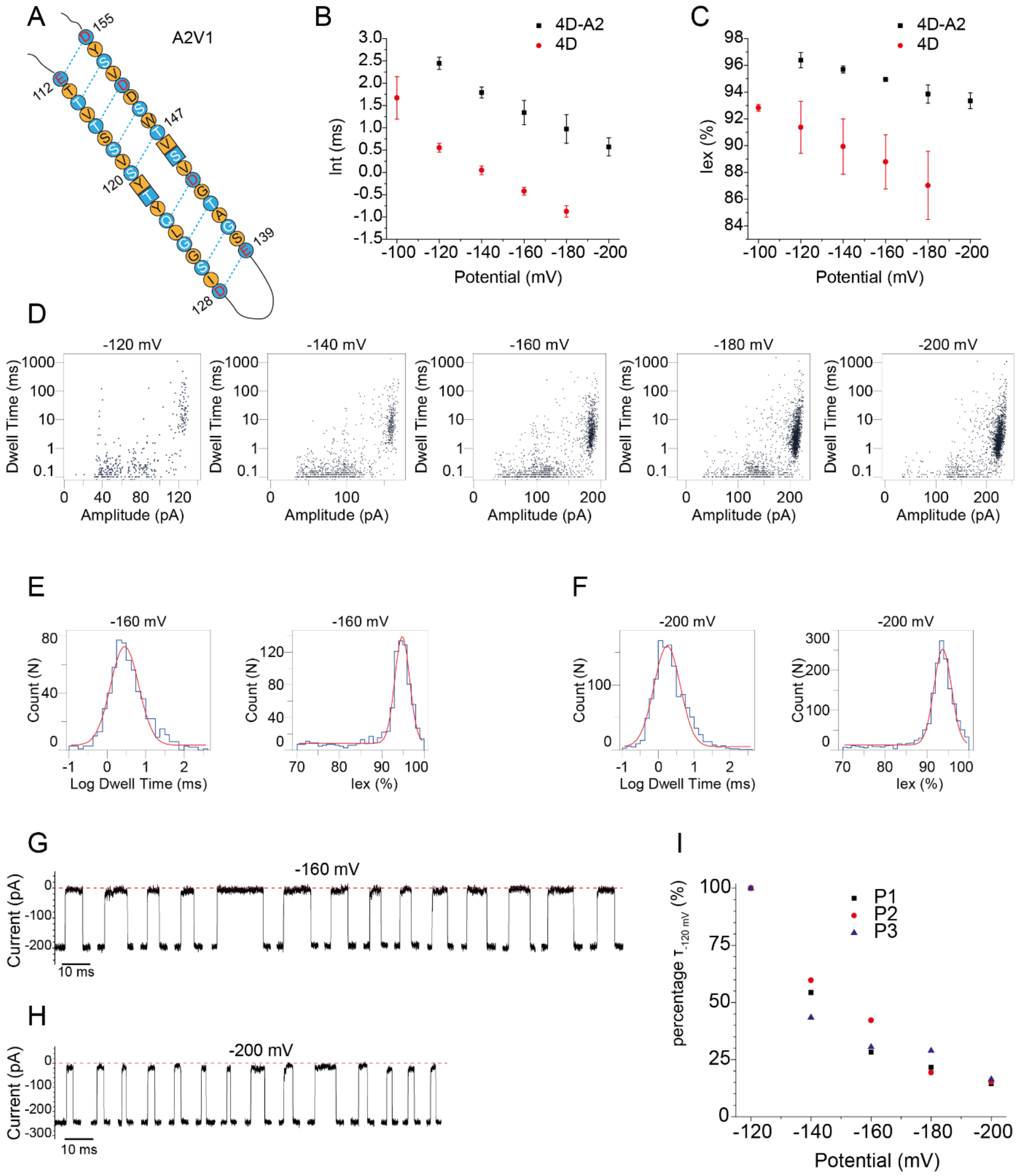

**Figure S12: Translocation of tzatziki across the A2V1 construct from the *cis* side (in $PBD_{11}PEO_8$).** A) Schematic representation of the extension within one monomer (only β-barrel domain depicted). Lumen-facing residues are shown in blue and membrane-facing residues are shown in orange. The residues present in the 4D pore are shown as circles and the newly introduced residues as squares. B-C) Natural logarithm of the dwell time (ln(τ)) (B) and $I_{ex}$% (C) dependence on the applied potential. Each data point represents the average from at least 3 nanopores and the corresponding standard deviation (SD). D) Scatter plots at the sampled potentials. E-F) Examples of log(dwell time) and $I_{ex}$% histograms obtained at -160 mV (E) and -200 mV (F). G-H) Examples of events at -160 mV (G) and -200 mV (H). I) Dwell time changes for individual pores expressed as percentage (%) of the dwell time at the lowest potential where translocation is observed (-120 mV). Recordings were performed in 1 M KCl, 15 mM Tris, pH 7.5, 50 kHz sampling rate and 10 kHz Bessel filter.

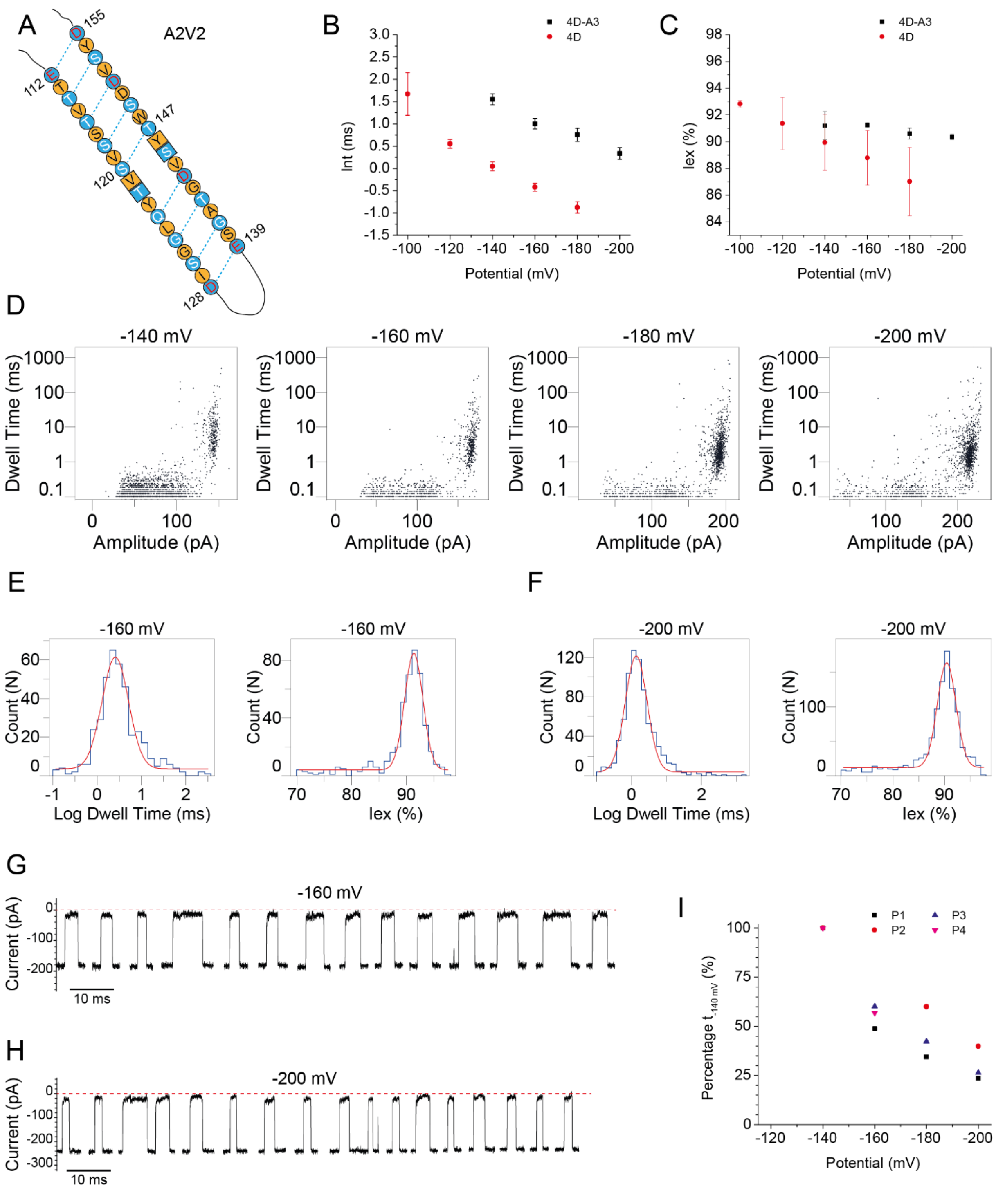

**Figure S13: Translocation of tzatziki across the A2V2 construct from the *cis* side (in $PBD_{11}PEO_8$).** A) Schematic representation of the extension within one monomer (only β-barrel domain depicted). Lumen-facing residues are shown in blue and membrane-facing residues are shown in orange. The residues present in the 4D pore are shown as circles and the newly introduced residues as squares. B-C) Natural logarithm of the dwell time (ln(τ)) (B) and $I_{ex}$% (C) dependence on the applied potential. Each data point represents the average from at least 3 nanopores and the corresponding standard deviation (SD). D) Scatter plots at the sampled potentials. E-F) Examples of log(dwell time) and $I_{ex}$% histograms obtained at -160 mV (E) and -200 mV (F). G-H) Examples of events at -160 mV (G) and -200 mV (H). I) Dwell time changes for individual pores expressed as percentage (%) of the dwell time at the lowest potential where translocation is observed (-140 mV). Recordings were performed in 1 M KCl, 15 mM Tris, pH 7.5, 50 kHz sampling rate and 10 kHz Bessel filter.

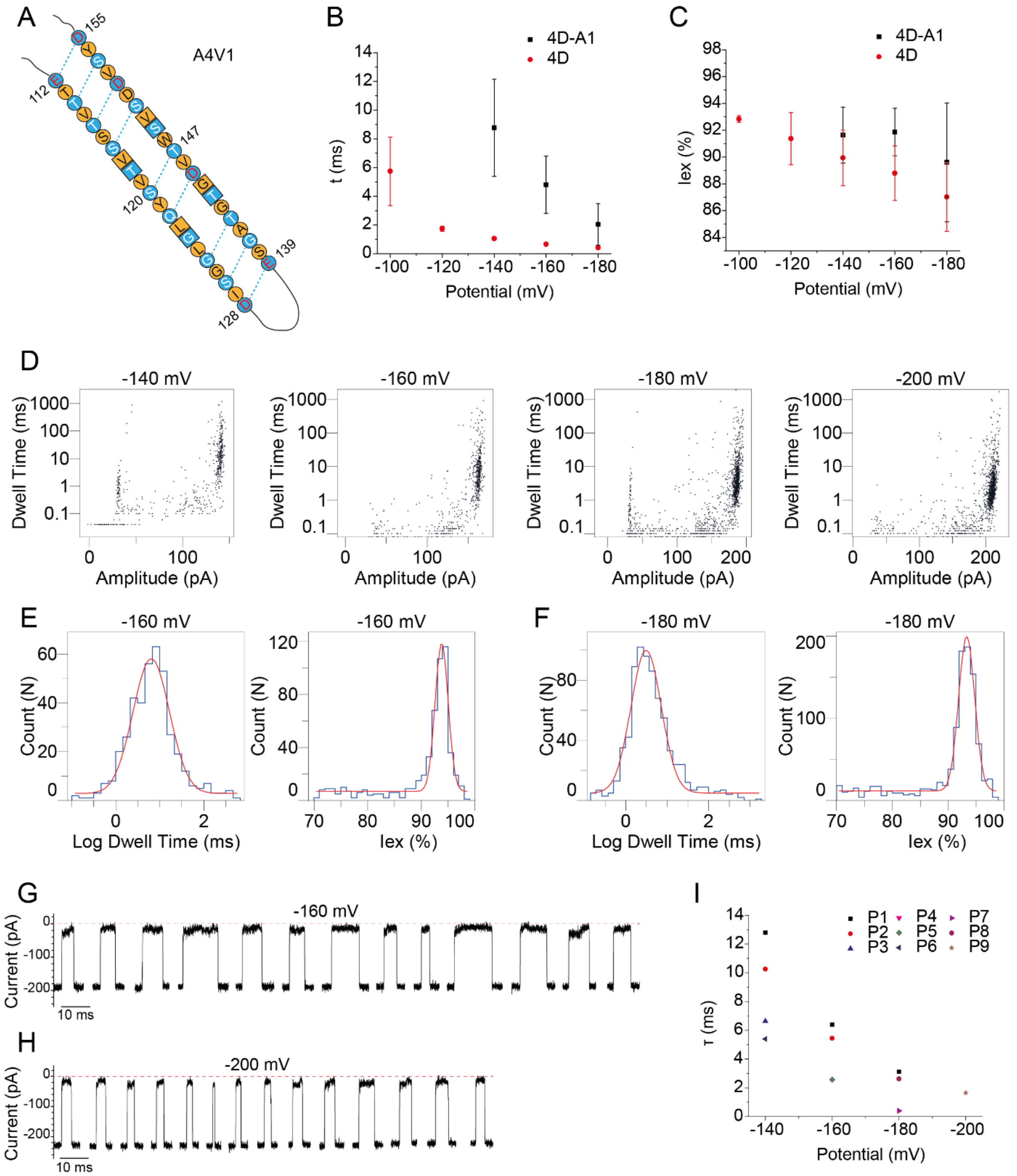

**Figure S14: Translocation of tzatziki across the A4V1 construct from the *cis* side (in PBD$_{11}$PEO$_8$).** A) Schematic representation of the extension within one monomer (only β-barrel domain depicted). Lumen-facing residues are shown in blue and membrane-facing residues are shown in orange. The residues present in the 4D pore are shown as circles and the newly introduced residues as squares. B-C) The dwell time (B) and $I_{ex}$% (C) dependence on the applied potential. Each data point represents the average from at least 3 nanopores (except for -200 mV, 1 nanopore) and the corresponding standard deviation (SD). D) Scatter plots at the sampled potentials. E-F) Examples of log(dwell time) and $I_{ex}$% histograms obtained at -160 mV (E) and -180 mV (F). G-H) Examples of events at -160 mV (G) and -180 mV (H). I) Dwell times from individual nanopores. Recordings were performed in 1 M KCl, 15 mM Tris, pH 7.5, 50 kHz sampling rate and 10 kHz Bessel filter.

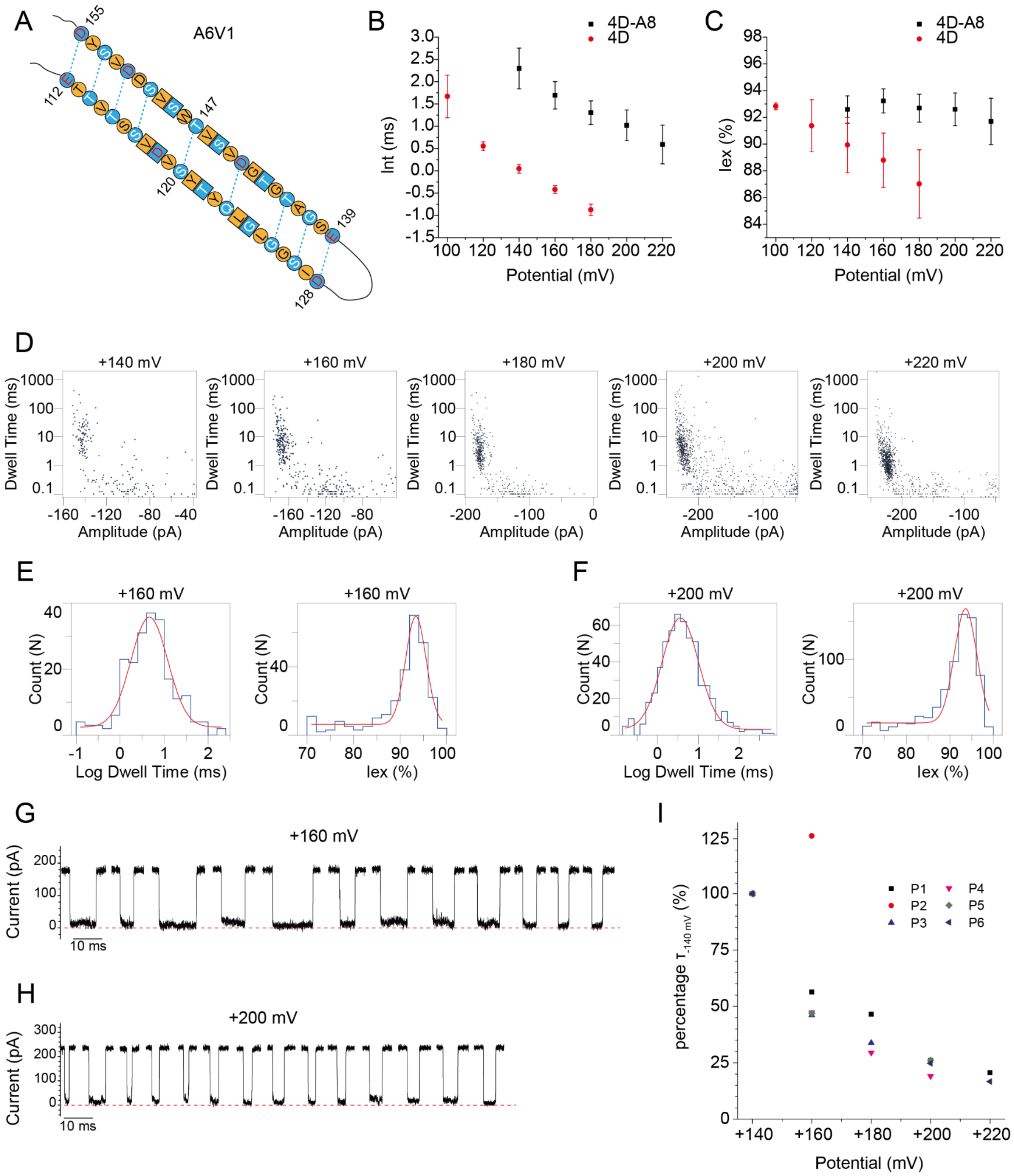

**Figure S15: Translocation of tzatziki across the A6V1 construct from the *trans* side (in $PBD_{22}PEO_{14}$).** A) Schematic representation of the extension within one monomer (only β-barrel domain depicted). Lumen-facing residues are shown in blue and membrane-facing residues are shown in orange. The residues present in the 4D pore are shown as circles and the newly introduced residues as squares. B-C) Natural logarithm of the dwell time (ln(τ)) (B) and $I_{ex}$% (C) dependence on the applied potential (negative for 4D pore, positive for 4D-A8). Each data point represents the average from at least 3 nanopores (except for +220 mV, 2 nanopores) and the corresponding standard deviation (SD). D) Scatter plots at the sampled potentials. E-F) Examples of log(dwell time) and $I_{ex}$% histograms obtained at -160 mV (E) and -200 mV (F). G-H) Examples of events at -160 mV (G) and -200 mV (H). I) Dwell time changes for individual pores expressed as percentage (%) of the dwell time at the lowest potential where translocation is observed (+140 mV). Recordings were performed in 1 M KCl, 15 mM Tris, pH 7.5, 50 kHz sampling rate and 10 kHz Bessel filter.
